# Dysregulated mammalian estrus cycle rescued by timed activation of VIP neurons in the circadian pacemaker and late afternoon light exposure

**DOI:** 10.1101/2023.01.14.524075

**Authors:** Anat Kahan, Gerard M. Coughlin, Máté Borsos, Bingni W. Brunton, Viviana Gradinaru

**Author notes:** Correspondence to V.G. and A.K.

## Abstract

Jet lag and shift work disrupt the menstrual cycle and decrease fertility. The circadian pacemaker, the suprachiasmatic nucleus (SCN), is known to modulate ovulation, but the mechanism is unclear. Here we explore this connection by tracking the dynamics of vasoactive intestinal peptide (VIP)-expressing neurons in the SCN in freely-behaving mice. We show that SCN^VIP^ activity is time-of-day- and sex-dependent, and estrous-state-dependent in late afternoon, gating downstream activation of GnRH neurons. Afternoon light, as well as specific activation of SCN^VIP^ neurons, rescues estrous cycle regularity and egg release in animals in altered light conditions, emphasizing the role of SCN^VIP^ neurons as a time-dependent light-responsive switch. Our results reveal the dynamic mechanism by which SCN^VIP^ neurons mediate light responses to regulate estrous states and demonstrate light-induced fertility rescue.

**One Sentence Summary:** Modulating and recording the activity of suprachiasmatic VIP neurons in freely behaving mice reveals their regulation of fertility by mediating the response to late afternoon light.

## Main

In the mammalian estrous cycle, fluctuations in sex hormone secretion occur with periods longer than a day, and these infradian rhythms are coordinated with the behavioral circadian rhythm to align the fertility window with social interaction and receptivity, thus maximizing reproduction (*1*). The critical event of ovulation is controlled by the hypothalamic-pituitary-gonadal (HPG) axis and, in mice, is synchronized with the dark phase every four or five days. In metestrus and diestrus, estrogen secretion from the ovary increases, followed by a luteinizing hormone (LH) surge in the late afternoon of proestrus, triggering ovulation ∼12 hours later. Hypothalamic neuronal populations such as gonadotropin-releasing hormone (GnRH) neurons in the medial preoptic area (MPA) (*2*), RF-amide-related peptide-3 (RFRP-3) neurons in the dorsomedial hypothalamus (DMH) (*3*) and kisspeptin (KISS1) neurons in the anteroventral periventricular nucleus (AVPV) (*4, 5*) and the arcuate nucleus (*6*) mediate estrogen feedback and are responsible for the LH surge.

The hypothalamic suprachiasmatic nucleus (SCN), the circadian rhythm pacemaker, is fundamental to ovulation occurrence. Several pieces of evidence suggest that signals from the SCN directly generate the LH surge. First, the SCN has neuronal projections to relevant hypothalamic regions, including the MPA (*7*), AVPV (*8*), and DMH (*8-10*). Second, SCN lesions cause a decrease in LH surge and estrous cycle regularity (*11-14*). Two circadian neuropeptides have been associated with the LH surge: arginine-vasopressin (AVP) and vasoactive intestinal polypeptide (VIP). Female mice deficient in VIP show reduced estrous-cycle regularity and fertility (*15*), and neurons expressing these peptides project directly or indirectly to GnRH (*16-18*), Kiss1 (*6*), and RFRP-3 (*10*) neurons. Specifically, VIP receptor type 2 (VPAC2, encoded by *Vipr2*) is present in ∼40% of GnRH neurons during proestrus, and VIP processes have been observed in apposition to GnRH neurons (*7, 19*). Based on bilateral differences in c-Fos expression in GnRH neurons in hamsters, it was suggested that circadian regulation of GnRH neurons is mediated by direct synaptic input to GnRH neurons (*20*). While VIP is known to excite GnRH neurons in brain slices, the estrous cycle regulation of this process is debated in the literature (*16, 18, 21*), and the infradian dynamics of VIP-expressing neurons throughout the estrous cycle are unknown.

Here, we show that light information controls estrous cycle regularity through SCN^VIP^ neurons. Our *in vivo* approach combines neuronal and physiological outcomes, which can be tracked long-term. Measuring the dynamics of SCN^VIP^ neurons across the estrous cycle using a calcium indicator with fiber photometry followed by advanced machine-learning-assisted signal analysis, we show that SCN^VIP^ neuronal activity is sex-specific and light-active, with increased activity in the late afternoon, especially when females are non-receptive. We observed a reduction in estrous cycle regularity when VPAC2 was downregulated in GnRH neurons, suggesting that this time-of-day information is gating the estrous cycle through GnRH neurons. Based on these observations, we hypothesized that afternoon light could rescue estrous cycle regularity under minimal dark conditions, as indeed we found, together with the rescue of egg release. Furthermore, this rescue could be mimicked by specific SCN^VIP^ activation in the afternoon, emphasizing the unique role of SCN^VIP^ neurons as a light switch. Together, this work establishes a solid mechanistic connection between light, SCN^VIP^ activity, estrous cycle, and fertility.

## Results

### Estrous cycle regularity in mice is significantly reduced by jet lag or ablation of SCN^VIP^ neurons

In humans, disruptions in the regular light-dark cycle (e.g., during jet lag) are associated with irregular menstrual cycling and decreased fertility. To confirm the significance of light regularity on the mouse estrous cycle, we performed a “jet lag” experiment in which we housed female C57 mice in 12:12 hour light:dark (LD) conditions, and every 4-5 days, advanced the light in the room by six hours. This pattern was previously found to dramatically decrease pregnancy success in mice (*22*), and reduce general fertility in humans (*23, 24*). Using cytology of daily vaginal smears over three weeks (4-5 cycles in non-jet-lagged animals), we found that this light pattern decreased the regularity of the estrous cycle in the mice, reducing the number of proestrus events from 4.6±0.2 to 2.1±0.3 (mean±sem) in three weeks. This effect was due mainly to increased time spent in estrus, from 7.0±0.6 to 9.1±0.9 days (Figure 1A-D).

**Figure 1:**
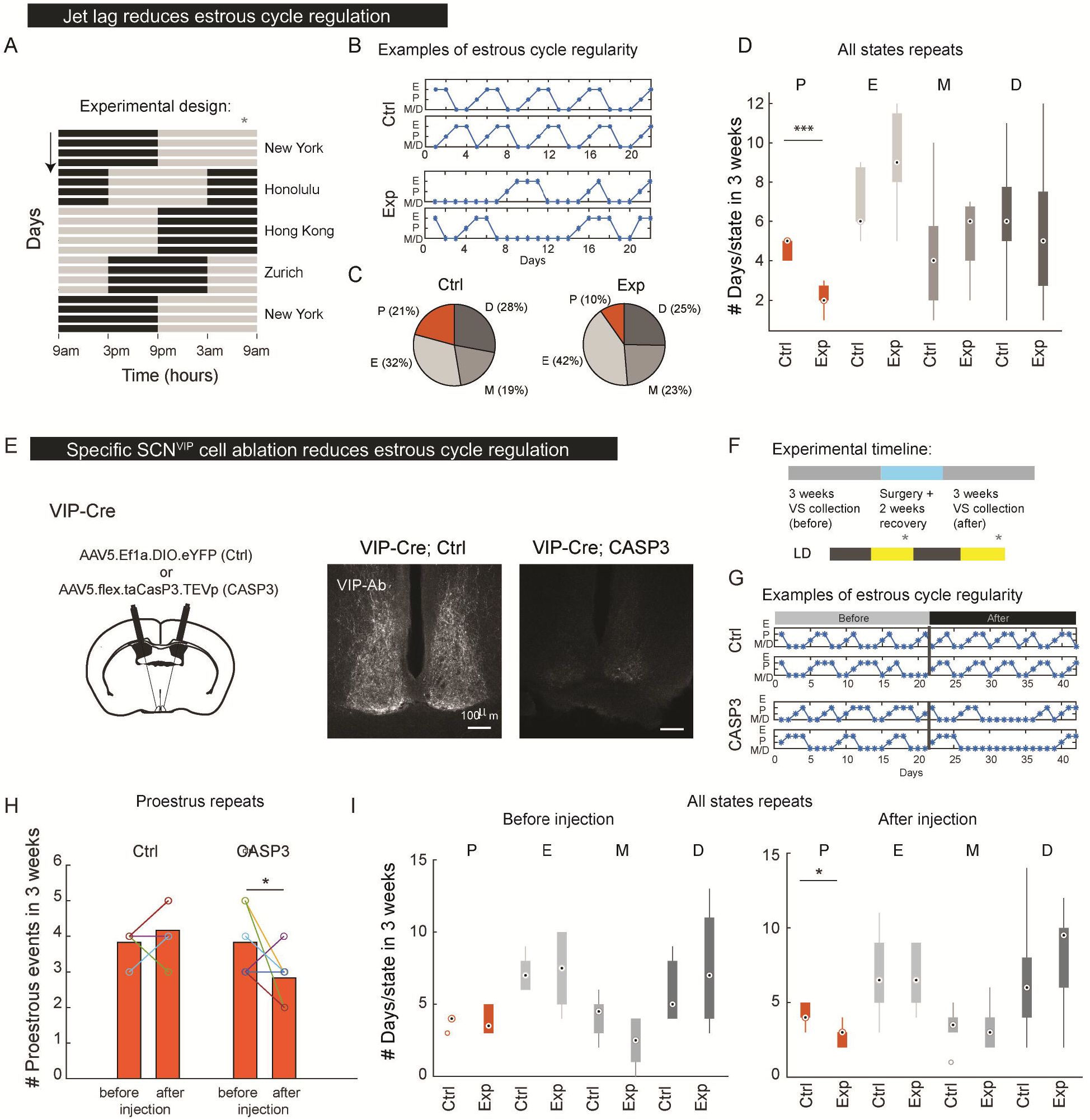
Estrous cycle regularity is significantly reduced by either jet lag or specific ablation of SCN^VIP^ neurons. (A) Jet lag experimental design, with light advanced shifts equivalent to flying from New York to Honolulu, four days later to Hong Kong, followed by Zurich and back to New York, all in three weeks. The gray asterisk indicates the time of vaginal smear collection. (B) Examples of vaginal smear traces of two control (top) vs. two jet-lagged (bottom) mice. P-proestrus, E-estrus, M-metestrus, D-Diestrus. (C) Distribution of estrous-cycle stages in C57 mouse females in jet-lagged vs. control periods (Ctrl: n=7, Jet lag: n=7). (D) Boxplot showing the number of days spent in each estrous stage in a three-week. (E) SCN^VIP^ ablation viral approach and VIP antibody staining in the SCN of a representative experiment. Experimental (CASP3, n=6) and control (Ctrl, n=6). (F) Female mice were placed under 12:12 LD conditions, and VS were collected in the late afternoon (gray asterisk) for three weeks, before and after virus injection, with two weeks of recovery after the surgery. (G) Examples of vaginal smear traces of two control (top) vs. two CASP3 injected (bottom) mice. The gray line separates before and after injection. (H) The number of proestrus events in three weeks, before and after injection of Cre-dependent CASP3 or control (Ctrl) virus. (I) Boxplot showing the number of days spent in each estrous stage in a three-week, before and after virus injection (*p<0.05, ***p<0.005; Nonparametric Kruskal–Wallis test. Two sample t-test when data was compared within two conditions and was normally distributed).

The SCN is one of the initial brain relays for retinal responses to light, and SCN^VIP^ neurons were previously identified as essential to the circadian response to light (*25-29*), forming monosynaptic connections to intrinsically photosensitive retinal ganglion cells (ipRGCs) (*27, 30*) and mediating the rapid response to light during the dark phase (*26*). We, therefore, tested the effect of specific ablation of SCN^VIP^ neurons on estrous cycle regularity in female mice kept under 12:12 conditions. We injected adeno-associated virus (AAV) to deliver Cre-dependent caspase-3 to the SCN of VIP-Cre mice, inducing targeted apoptosis of SCN^VIP^ neurons (Figure 1E). We found that ablation significantly reduced the number of proestrus repetitions, from 4.3±0.4 to 2.8±0.3 in three weeks, significantly lower than the control group (Figure 1F-H). These results support a specific role for SCN^VIP^ neurons in connecting light stimulus with estrous cycle regulation.

### SCN^VIP^ activity is sex-specific and time-of-day-dependent

To characterize the temporal dynamics of SCN^VIP^ neuron activity, we used fiber photometry (FP) to record bulk neuronal activity in males and females using a genetically encoded calcium indicator, GCaMP6s, using VIP-Cre x GCaMP6s mice, implanted with two 400 µm optical fibers. To verify the fiber location, we used light-guided-sectioning (LiGS) (*25*), a 3D histological method that we developed to determine the precise position of optical implants within deep and small brain nuclei, such as the SCN (Figure 2A). First, we tested the responsiveness of the calcium signal to the light phase transitions, zeitgeber time (ZT) 11 to 13 and ZT23.5 to 0.5. Using dF/F and event-rate analyses, we found that SCN^VIP^ activity is synchronized with the light change (Figure S1). Having thus confirmed light sensitivity at the light phase transitions, we recorded bulk GCaMP6s activity for 10 minutes each hour over 24 hours (Figure 2B). We determined the estrous stage of cycling females by vaginal smear cytology over multiple continuous days until at least three cycles were recorded, and compared their SCN^VIP^ activity to that of males and ovariectomized (OVX) females, recorded for at least three days each. All animals showed circadian fluctuations, with reduced SCN^VIP^ activity during the dark hours of ZT12-24, based on both event rates (Figure 2C) and dF/F (Figure S2A-B). Females had higher integrated dF/F than males during the light phase (8.7±0.8 and 3.5±0.5 a.u., females and males, respectively, Figure S2A-C), but we did not identify significant differences in bulk activity between estrous stages or between intact and ovariectomized females (Figure 2D-E). We observed significantly increased event rates during the light phase for all estrous stages (0.36±0.04 vs. 0±0 events/minute, light vs. dark, averaged over estrous stages) and increased dF/F values for some (Figure S2E). Interestingly, we noticed that both event rate and dF/F exhibited parabolic activity during the light phase (Figure 2E and Figure S2F, averaged R^2^=0.8±0.07 mean±sem, ZT2 to 11), decreasing to an average minimum at ZT5 and then increasing again to a maximum at ZT11. By comparing specific time windows, we observed a trend of higher event rates in M/D compared to E during the late afternoon (ZT11-12), but none of the comparisons showed a significant difference between females in different estrous stages or OVX (Figure S2G). Based on this analysis, FP data suggest that the main role of SCN^VIP^ neuronal activity, which is sex-dependent, is a permissive circadian switch aligned with light information to convey the best timing to ensure fertility.

**Figure 2:**
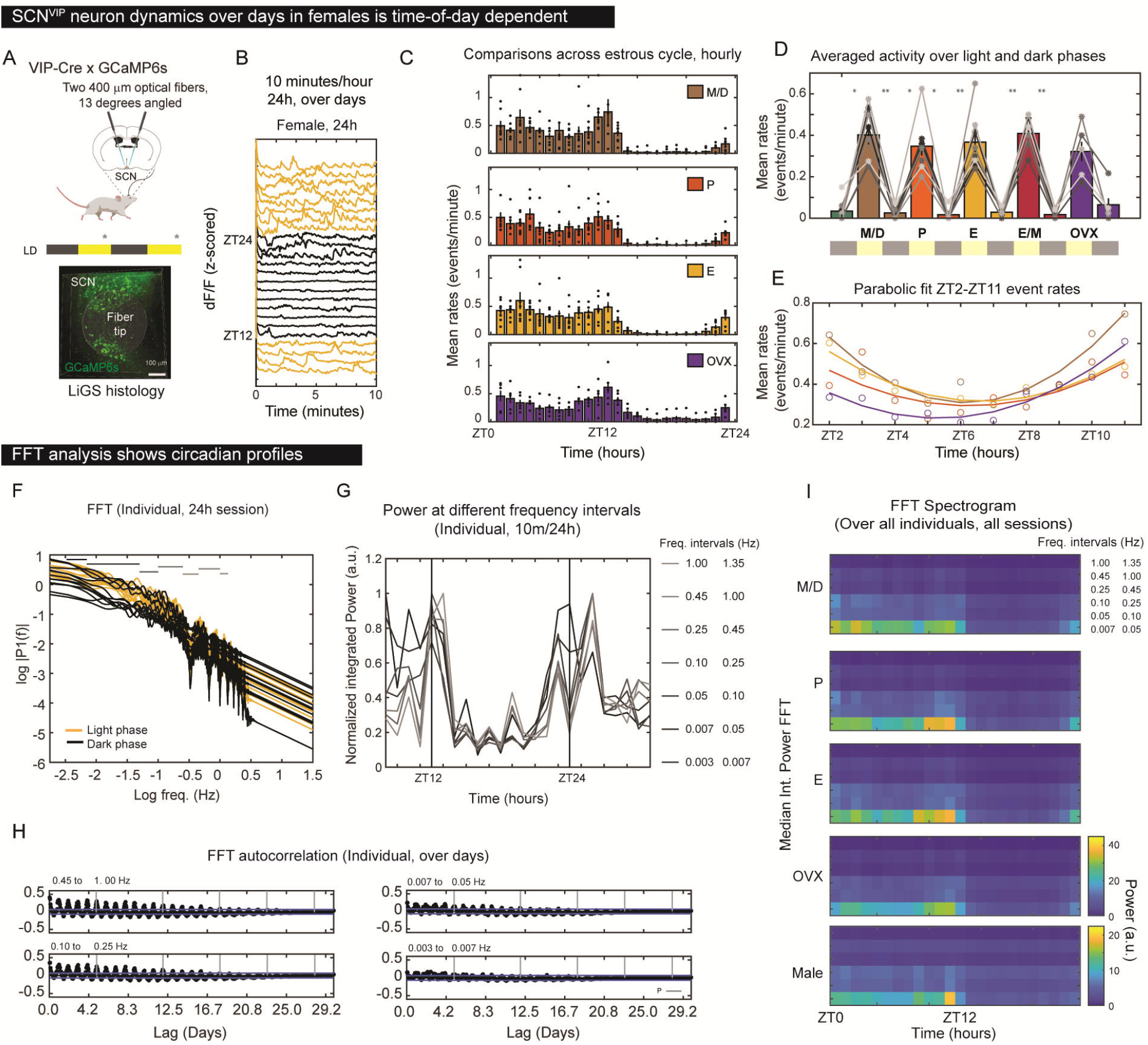
Temporal changes in SCN^VIP^ activity show circadian patterns of event rate and FFT spectrogram. (A) The experimental setup: VIP-Cre crossed to Ai162-reporter line to express GCaMP6s. The fiber location is validated with light-guided-sectioning (LiGS) histology (*25*). Mice were kept under 12:12 LD conditions. VS were taken in the late afternoon (gray asterisk) (B) A representative example of 10-minute-per-hour recording, over 24 hours, from one female. (C) Event rates over 24h of recording across estrous states (n females=8, n OVX=5, each state is represented at least three times in each female). (D) Event rates averaged over dark and light phases. For clarity, significance is marked only between adjacent periods. Nonparametric Kruskal-Wallis test, followed by Tukey’s correction (D) (*p<0.05; **<0.01). (E) Parabolic fits to event rates during ZT2-11, showing minimum at ZT5 and maximum at ZT11. (F-J) FFT analysis (See Figure S3 and Methods for details). (F) FFT power vs. frequency for one session of one animal (logarithmic scale). Each line represents 10 minutes of recording during the dark phase (black) and the light phase (yellow). The seven gray lines represent the integrated frequency intervals: 0.003 to 0.007, 0.007 to 0.05, 0.05 to 0.1, 0.1 to 0.25, 0.25 to 0.45, 0.45 to 1.0, and 1.0 to 1.35 Hz, respectively. (G) Integrated FFT powers of the frequency intervals shown in (F) over 24h, showing one cycle of light-dark response. (H) FFT autocorrelation over days for four of the frequency intervals, showing circadian modulations. The interval of 0.003 to 0.007 Hz has reduced circadian phenotype and therefore removed from further analysis. (I) Averaged FFT spectrograms for all recorded sessions by sex and hormonal state, showing clear 12:12 LD contributions.

FP is a bulk measurement. Reasoning that subpopulations of VIP neurons within the SCN may have different activity properties (*31*), we applied fast Fourier transform (FFT) analysis, a method to transform time-series data from the time to the frequency domain, and therefore separate the frequencies of time-variant oscillators (*32*). We applied FFT to each 10 minutes recording-period, for each 24h session and identified seven frequency ranges (from 0.003 to 1.35 Hz) with distinct log-log profiles (Figure 2F). Integrating the power spectra over these frequency ranges showed light-dark rhythmicity (Figure 2G), and autocorrelation over the multiple days of recording revealed circadian phase in all frequency ranges (Figure 2H), with low amplitude at the lowest range (0.0033 to 0.007 Hz), which therefore was not included in the following analyses. Averaging the integrated power spectra of individual animals across experimental groups showed a clear difference in the amplitutes of the low-frequency SCN^VIP^ contributions between males and females (Figure 2I, note the different color bar range), all showing clear circadian profile, time-of-day dependent.

### Late-afternoon SCN^VIP^ activity correlates with estrous state

Next, we aim to quantify the differences in the FFT spectrograms, focusing on the uniqueness of the late afternoon, which is the time frame of the LH surge. We, therefore, looked at differences in low-frequency FFT signal between estrous states. FFT histograms of males was distinct from OVX females throughout the light period (Figure 3A). By contrast, we saw a time-of-day specific difference in FFT histograms between cycling females on non-receptive (M/D) and receptive (E) days, with a separation at ZT9, but not ZT4 (Figure 3B). To further investigate this critical window, we tested whether the qualitative differences in FFT histograms we observed were sufficient for a machine-learning-based classifier to discriminate between states. We applied a leave-one-out cross-validation (LOOCV) approach to the FFT spectrograms, including frequencies up to 1 Hz, as extracted from individual 10 minutes/hour FP daily trials of males, OVX females, and cycling females. In each round, an even number of trials from two conditions was used to train the classifier, which was then tested on an additional trial not included in the training set (using “Support vector machine” algorithm, SVM, see Methods for details). The overall score reflects the prediction success of the classifier over all rounds. As expected, the classifier was able to distinguish between males and females, with an average accuracy score of 74%. Interestingly, the classifier could also distinguish OVX females from both males and other females (Figure 3C), reflecting the increased sensitivity of the FFT approach compared to simple event detection.

**Figure 3:**
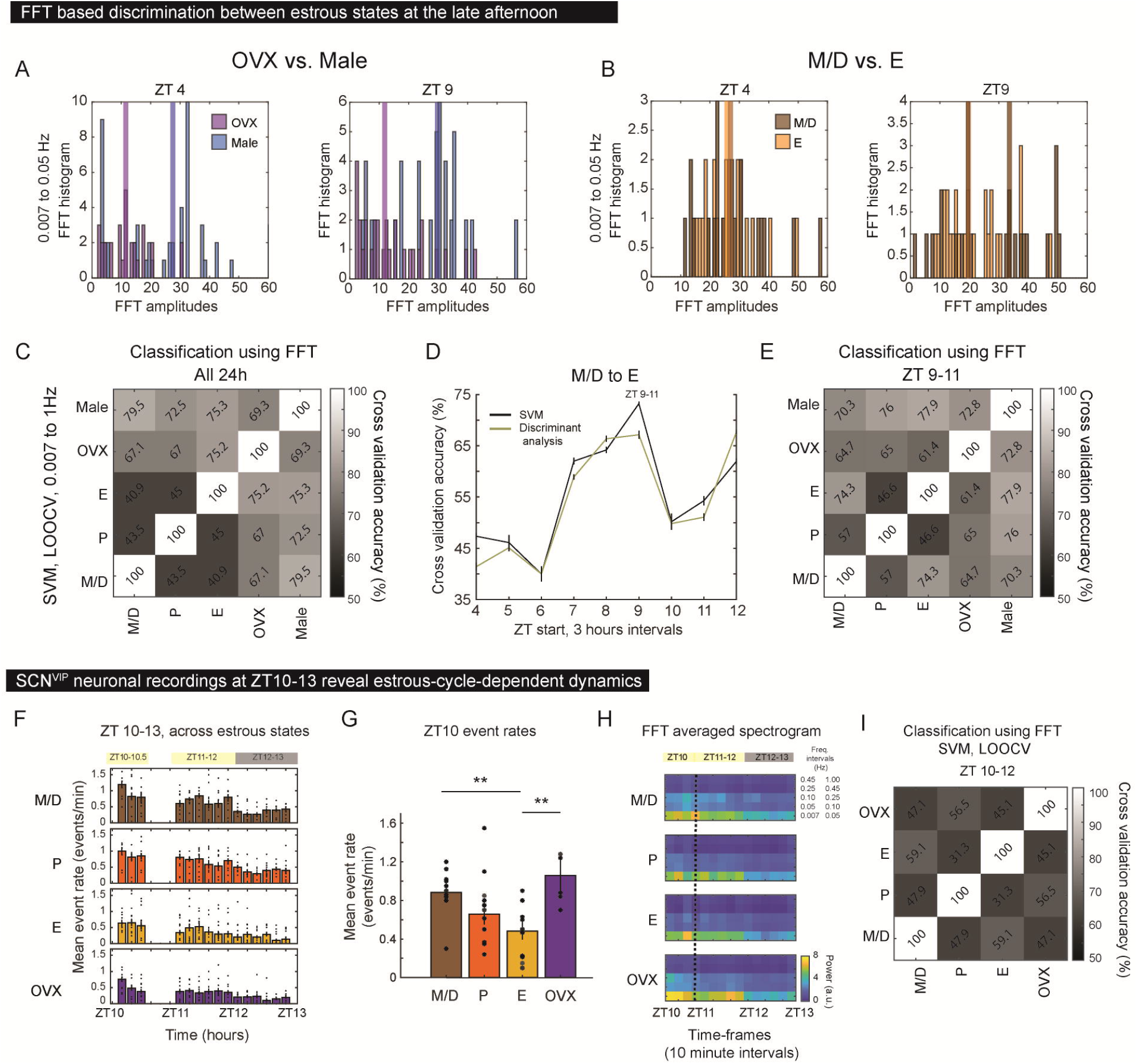
Estrous-cycle is discriminated in late afternoon with FFT classification and additional wide temporal window afternoon recording of SCN^VIP^ FP signal. (A) FFT histograms at ZT4 and ZT9 of the frequency range 0.007 to 0.05Hz, comparing OVX females and males. Thick lines show the histogram medians. P=0.00003 and P=0.0002 at ZT 4 and 9, respectively, two-sample Kolmogorov-Smirnov test. (B) Same as (A), for M/D vs. E. P=0.88 and P=0.02 at ZT 4 and 9, respectively, two-sample Kolmogorov-Smirnov test. (C) Cross-validated accuracy with support vector machine (SVM) classifier for discriminating sex and hormonal state prediction ability, based on full FFT spectrograms (24h), averaged over five double-sided iterations for OVX, males, and females on different estrous cycle days. Males: n=6; OVX: n=5, Females: n=6; Chance level is 50%. see Figure S4 for a comparison with “Discriminate analysis” classifier. (D) Cross validation accuracity score for M/D vs. E, using FFT at intervals of three hours, comparing SVM (black) with “Discriminate analysis” algorithm (green), both showing a peak at ZT9-11. (E) FFT-based classification, based on FFT spectrograms at ZT9-11, averaged over five double-sided iterations. (F-I) SCN^VIP^ neuronal dynamics at ZT10-13. (F) Event rates during ZT10 to 13, divided to 10 minutes intervals, of cycling and OVX females. (G) Averaged event rates at ZT10-10.5. Nonparametric Kruskal-Wallis test, followed by Tukey’s corrections (**p < 0.01). (H) Averaged FFT spectrograms across all states. (I) Cross-validation accuracy of SVM classifiers of the discrimination prediction ability across estrous cycle days.

We used the cross-validation score to test whether within the total 24h of activity there is a relevant time period for estrous cycle classification, focusing on the ability to distinguish between before ovulation (M/D) and immediately after ovulation (E). Indeed, the prediction ability between M/D and E increased at the late afternoon with a three hours time window, which increased the chance level at ZT7-10, with maximal discrimination ability at ZT9-11 (Figure 3D). Using only this critical window of ZT 9-11 enabled the classifier to predict females’ receptivity status, E vs. D/M, with an accuracy of 74% (Figure 3F, supported by using an alternative algorithm, Figure S4), and allows above change classification between before and after ovulation (M/D vs. P).

Reasoning that we might have missed transient changes in SCN^VIP^ signal in the critical window by recording only 10 minutes per hour, we repeated our FP experiment with continuous recording from ZT 10-10.5, the approximate time of the LH surge during proestrus, and from ZT 11-13. We observed a significant decrease in event rate at ZT10 when females were in E, compared to M/D (Figure 3H), in line with the trend we observed with 10 minutes per hour sampling (Figure 2D). We next applied FFT analysis, binning the continuous recordings into ten-minute intervals (Figure 3I), and found that the classifier could now discriminate not only between females in M/D and E (59.8±0.3%, mean±sem, over ten iterations) but also between OVX females and females in P or E, with prediction success over 65% (Figure 3J).

Since the SCN is estrogen-sensitive (*33-35*), we decided to test whether hormonal stimulation of proestrus in OVX females would recover a receptive pattern of SCN^VIP^ activity. However, hormonal induction with estrogen (OVX+E) followed by progesterone (OVX+E+P) did not significantly alter SCN^VIP^ activity at ZT10 (Figure S5B), suggesting that the ovariectomy had already caused irreversible changes in the SCN (*25*). Together, our FP results show a late afternoon decrease in SCN^VIP^ activity during the receptive period, possibly indicating a mechanism to inhibit the downstream HPG target just before and after ovulation.

### Loss of VIP receptors on GnRH neurons disrupts estrous cycle regularity

Our data suggest that SCN^VIP^ neurons act as a permissive circadian switch, integrating light information to inform optimal fertility timing. We next sought to identify the target of this switch. Prior slice electrophysiology showed that VIP increases the activity of GnRH neurons in a time-of-day dependent manner (*16, 18*). While these results are suggestive of a direct effect, it could also be indirect, affecting local SCN interactions, such as to AVP neurons, which further project to Kiss1 neurons (*36, 37*), the pulse generator for GnRH neurons (*38*), or via ovarian circadian control (*12, 39*). To test whether SCN^VIP^ neurons transmit light information directly to GnRH neurons, we used Clustered Regularly Interspaced Short Palindromic Repeats (CRISPR) genetic editing to disrupt either VPAC2 or Kiss1 receptor (KISS1R) expression specifically in MPA^GnRH^ neurons that project to the median eminence (ME) by AAVretro injection into the ME (*40*) (Figure 4A). We verified MPA^GnRH^ transduction and receptor knockdown by immunostaining (Figure 4B-D), which showed a significant physiological effect despite low transduction efficiency. Female mice were kept in 12:12 LD conditions, and their estrous cycles tracked for three weeks prior to and after CRISPR editing (following a two-week recovery period after surgery). While the control group exhibited a regular estrous cycle, with an average frequency of five proestrus events in three weeks, knockdown of either *kiss1r* or *vipr2* resulted in a significant reduction in proestrus events (from 4.2±0.3 to 2.7±0.3, p=0.0 for *kiss1r* and from 4.8±0.3 to 2.8±0.3, p=0.007, for *vipr2*, Figure 4E). Similar to what we observed with SCN^VIP^ ablation, females with disrupted VPAC2 expression also spent significantly more time in estrus (Figure 4F). These results suggest that SCN^VIP^ act directly on GnRH neurons through VIP release, to further regulate the estrous cycle.

**Figure 4:**
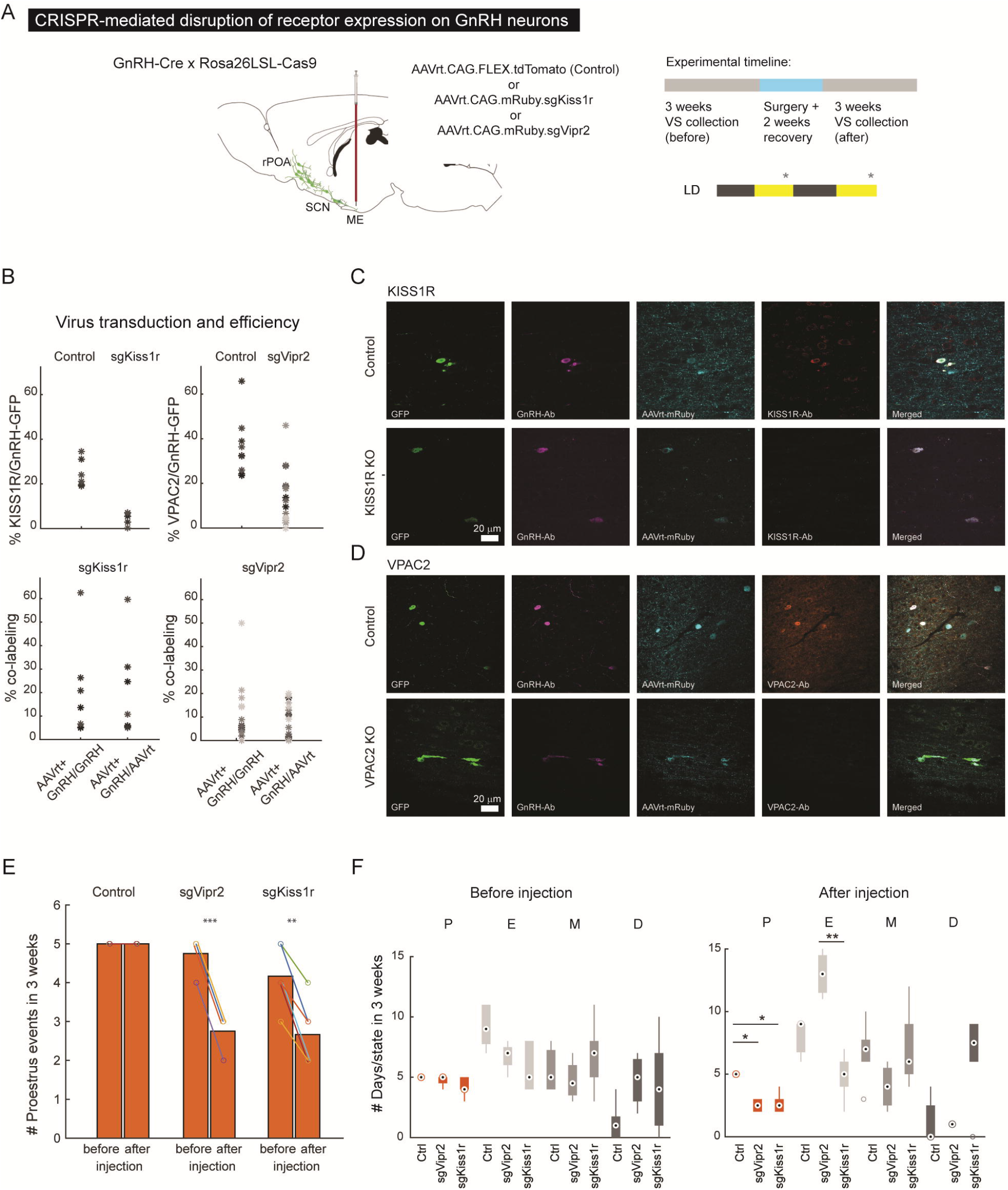
CRISPR-Cas9 mediated disruption of VPAC2 and KISS1R on GnRH neurons reduces estrous cycle regularity. (A) Experimental design: viral injection to the ME with AAVretro, to cover all GnRH neurons which project to the ME. VSs were taken for three weeks, before and after virus injection. (B) Transduction efficiency in experimental mice for KISS1R (i) and VPAC2 (ii), three samples from each animal, represented by different shades of gray. (C) Representative examples of MPA histology, showing KISS1R-Ab expression (orange) overlapping with GnRH neurons (Cas9 in green, GnRH-Ab in purple) in control mice (top), but not in experimental mice (bottom), which show AAVretro expression (cyan). (D) Same as (C), for VPAC2. (E) The number of proestrus events identified in three weeks, before and after virus injection in control (n=5), sgKiss1r (n=6), and sgVipr2 (n=4) mice. (F) Estrous states appearance the period of three weeks, comparing all conditions. Estrous state distributions by percentage are available in Figure S6. (*p<0.05; **<0.01, ***<0.005; Kruskal–Wallis test, Tukey’s correction for multi comparisons).

### Afternoon light rescues estrous cycle regularity and egg release under limited light conditions

The time-of-day-dependent activity of SCN^VIP^ neurons that we observed inspired us to test whether light exposure in the afternoon sensitive time window would be sufficient to rescue circadian-disrupted fertility *in vivo*. To limit light exposure while minimally affecting locomotor activity, we group-housed females in constant-dark conditions (DD) with a 30-minute pulse of light at circadian time (CT) 0 (DD+CT0_0.5L_) (Figure S7). We then tested different light stimulus regimes sequentially for three weeks each while tracking estrous stages: LD, DD, DD+CT0_0.5L_, and two conditions with an additional hour of light, the first at CT10 (DD+CT0_0.5L_+CT10_L_) and the last at CT4 (DD+CT0_0.5L_+CT4_L_, Figure 5A). During the LD period, we observed an average of 4.2±0.5 proestrus events in three weeks, indicative of a normal estrous cycle. DD and DD+CT0_0.5L_ conditions reduced this to 2.8±0.8 and 1.8±0.3 proestrus events in three weeks, respectively. Notably, one hour of light at CT10, but not CT4, was sufficient to rescue the regularity of the estrous cycle, with 4.8±0.3 proestrus events in three weeks and regular distribution of estrous states (Figure 5B-C). To directly assay fertility outcome, we also measured the number of eggs released to the oviduct on the day following ovulation, identified by the transition from proestrus to estrus. We avoided using pregnancy success as a fertility readout since pregnancy itself might involve circadian clock cues (*22*). Compared to age-matched controls in 12:12 LD conditions, females in DD+CT0_0.5L_+CT10_L_ conditions released approximately twice as many eggs as those in DD+CT0_0.5L_ conditions (Figure 5D). Together, these results show that ambient light in the late afternoon is critical for both estrous cycle regularity and increased egg release.

**Figure 5:**
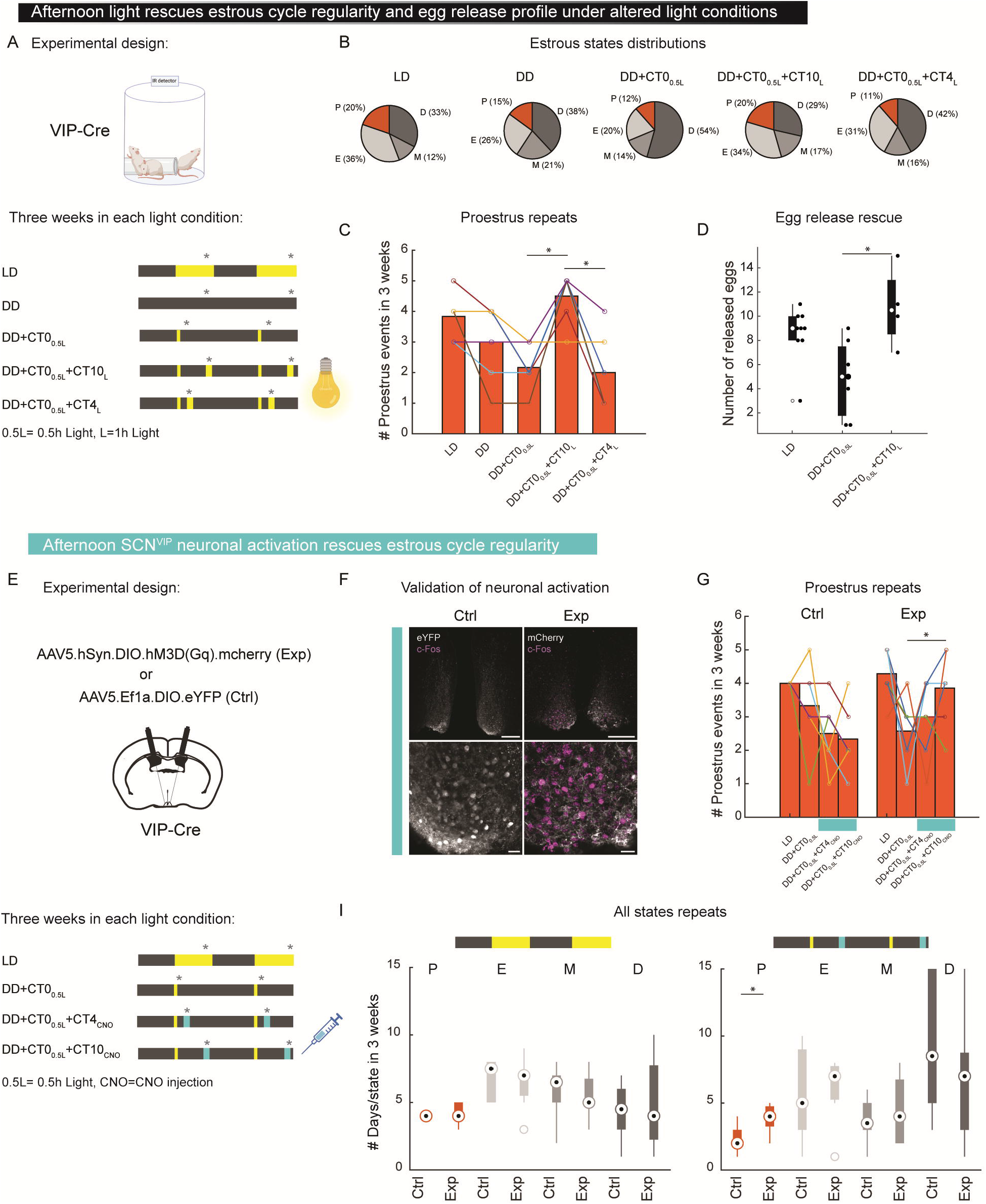
Under altered light conditions, afternoon light rescues estrous cycle regularity and egg release profile, and specific SCN^VIP^ neuron activation with excitatory DREADD rescue estrous cycle regularity. (A) Light cycle manipulation experimental design. Yellow: light. Dark gray: dark. Gray asterisks indicate the time of vaginal smear collection, aiming to minimize light exposure during the dark, even to ambient red light. CT0_0.5L_ refers to 0.5h of light at CT0. CT4_L_, or CT10_L_ refers to 1h of light at CT4 or 10. CT0 is the activity offset in nocturnal animals (see LMA in Figure S7) (B) Estrous stage distribution under the five light conditions (VIP-Cre, n=4), P-proestrus, E-estrus, M/D-metestrus, or diestrus. (C) The number of proestrus events in three weeks (mean). (D) The number of released eggs in three light conditions (n= 9, 7, 4 per condition, respectively). Boxplot showing median (white dot), 25^th^ (bottom), and 75^th^ (top) percentile. (E) Chemogenetic experimental design. VIP-Cre females were injected with Gq expressing virus (Exp, n=7) or eYFP (Ctrl, n=6), and put under four conditions, LD, DD+CT0_0.5L_, DD+CT0_0.5L_+CT4_CNO_, and DD+CT0_0.5L_+CT10_CNO_ (CT0_L0.5_, CT4_CNO_, and CT10_CNO_ correspond to 0.5h light at CT0, CNO injection at CT4 and CT10, 1mg/kg, cyan). (F) *Post hoc* histology, showing virus expression, mCherry (gray, Exp) and eYFP (gray, Ctrl), with c-Fos staining (purple), ∼1h after CNO application. (G) The number of proestrus events in three weeks, under the three conditions shown in E. (I) Overall appearance of the estrous states in the LD condition and in the DD+CT0_0.5L_+CT10_CNO_ condition. Estrous state distributions by percentage are available in Figure S8. Nonparametric Kruskal-Wallis test, followed by Bonferroni corrections (*p < 0.05).

### Activation of SCN^VIP^ neurons in the afternoon rescues estrous cycle regularity

Based on the ability of afternoon light to rescue estrous cycle regulation under limited light conditions, we hypothesized that specific SCN^VIP^ activation could do the same. We first tried an optogenetic approach in VIP-Cre x ChR2 mice using a similar temporal activation paradigm (Figure S9A). ChR2 activation at CT10 rescued proestrus frequency in half of the experimental cohort (VIP-Cre x ChR2, 4 females out of 8), compared to one VIP-Cre x ChR2 animal in the control group (no excitation) (Figure S9B). While we verified fiber localization using *post hoc* LiGS histology and activation of SCN^VIP^ neurons by c-Fos labeling under the fiber (Figure S9C), it is possible that the variable results were due to differences in fiber localization affecting the distribution of the light relative to the core of the SCN, heterogeneous distribution of neuronal subpopulations, or asymmetric connectivity of the SCN to the ovary (*41*).

While both optogenetic and chemogenetic methods have been used previously for SCN^VIP^ neurons (*26, 31, 42, 43*), chemogenetic approaches produce longer time scales of activation, which might better represent the natural activation of SCN^VIP^ neurons. We therefore used a single injection of clozapine N-oxide (CNO), a synthetic drug, to induce calcium flux through an excitatory Designer Receptor Exclusively Activated by Designer Drugs (DREADD), hM3Dq, delivered to the SCN by AAV injection (Figure 5E). We verified CNO activation *post hoc* by c-Fos expression (Figure 5F). Again, we tested the estrous cycle regularity of experimental and control (injected with AAV not carrying hM3Dq) groups over four sequential conditions (three weeks each): LD, DD+CT0_0.5L_, and DD+CT0_0.5L_ with CNO injection at either CT4 or CT10 (Figure 5E). We found that activation of SCN^VIP^ neurons in the critical time window at CT10, but not earlier in the day, rescued the regularity of the estrous cycle (Figure 5G-I).

## Discussion

Here we novelty show that either jet lag or specific ablation of SCN^VIP^ neurons significantly reduces estrous cycle regularity in mice. By measuring bulk calcium dynamics of SCN^VIP^ neurons over sequential days, we found that SCN^VIP^ neurons are primarily light-entrained and that their activity levels cycle over the light period. Using FFT analysis and advanced machine learning (ML)-based discrimination, we identified significant differences in activity patterns between males, naturally cycling, and ovariectomized females. In addition, we identified a difference in SCN^VIP^ activity at the peak time of ZT10 during receptive days. While ML has been used to identify sex differences in biological samples or behaviors (*44-47*) and estrous cycle stages from vaginal smears (*48-50*), this is, to our knowledge, the first demonstration of its use to identify estrous cycle states based on neuronal activity. Hypothesizing that SCN^VIP^ activity at ZT10 acts as a permissive signal to convey the best timing to ensure fertility, we found that SCN^VIP^ excitation either through the retinohypothalamic tract via light or via chemogenetically-specific activation at this unique feedback timepoint of ZT10 rescued estrous cycle regularity. To our surprise, afternoon light also rescued the number of eggs released in limited-light conditions, indicating a significant improvement in fertility. While the collection of released eggs is not commonly performed in naturally cycling mice, we were able to do so due to our close monitoring of the estrous cycle, thereby avoiding confounding effects of the exogenous hormones required to induce superovulation (*51*).

While SCN^VIP^ neuronal activity is known to participate in the control of the HPG axis, here we show for the first time, to our knowledge, that it is sex and ovarian-dependent *in vivo*. In the future, it will be of interest to determine the mechanism of hormonal control. In addition to a previously-reported dependency on estrogen (*25, 33-35*), we suspect progesterone is also involved, based on hypothalamic single-cell RNAseq data suggesting that VIP-targeted neurons co-express progesterone receptor (PR) mRNA (*52*). While we attempted to test this dependency *in vivo* using OVX females, we did not observe a recovery of receptive-like activity in hormonal stimulation paradigms; the long period following ovariectomy without hormonal replacement may have caused a loss of plasticity that brief hormonal treatment could not rescue.

The decreased ovulation we observed in a minimal light schedule aligns with previous reports of reduced estrous cycle regularity in response to altered LD (*53*) and DD schedules (*54, 55*). A role for VIP was first established by showing that a full-body VIP-KO female mouse had reduced estrous cycle regularity (*15*) and previous studies have established VIP-GnRH interaction, showing that VIP excites GnRH neurons in brain slices (*16, 18*) and reporting anatomical connectivity (*7, 17, 19*). Here, using new technologies, including opto- and chemogenetics, CRISPR gene editing and viral delivery, we were able to pinpoint the physiological effect on fertility to VIP neurons located in the SCN and downstream VIP receptors on GnRH neurons in the MPA. Note that other VIPergic populations outside the SCN might also signal to GnRH neurons through VPAC2 (*56*). In addition, VIP could also act through other paths, including local suprachiasmatic stimulation of AVP neurons, which are known to regulate Kiss1 neurons, or via peripheral stimulation of the ovary (*12, 41*). Consistent with the former, we observed estrous cycle disruption by knockdown of KISS1R in MPA^GnRH^ neurons. We also note that SCN^VIP^ neurons are not homogenous in at least two aspects. First, SCN^VIP^ neurons release two neuropeptides, VIP and GABA (*57*). We examined the role of VIP because it was shown that the calcium activity of SCN^VIP^ neurons correlates with VIP release (*26*), but we cannot rule out an additional role for GABA. Second, recent studies have functionally (*31*) and genetically (*27, 58*) defined two subpopulations of SCN^VIP^ neurons. It will be interesting to address these aspects in the future.

Based on our results, we suggest the following model: circadian oscillation of SCN^VIP^ neuronal activity gives a late-afternoon permissive input, with increased event rate during metestrus and diestrus, to downstream targets such as MPA^GnRH^ neurons, mediated by direct VIP signaling to GnRH neurons, as well as to other hypothalamic cell populations, such as SCN^AVP^ and possibly to the ovaries. SCN^AVP^ neurons in turn, signal to Kiss1 neurons, which, together with the direct SCN^VIP^ signal, ensure increased GnRH activity to drive the LH surge during proestrous. Following the LH surge, estrogen levels drop and progesterone levels rise, causing a decrease in SCN^VIP^ neuronal activity in this critical window of the following day (estrus). We suggest that this reduction in activity functions to prevent the reoccurrence of ovulation, possibly via reduced stimulation of GnRH neurons and increased activity of DMH^RFRP-3^ neurons (*10*).

Our work here in mice offers a model of a combination between circuitry and dynamics for studying the HPG axis and its neuronal control by the circadian rhythm pacemaker. This approach may be particularly useful in an era of circadian disruption due to artificial light and with increasing evidence that shifted working hours are associated with human reproductive disorders (*59*). In particular, our observation of increased egg release in response to targeted light intervention may inspire potential interventions for women who struggle with infertility due to shifted working hours or frequent jet lag (*60-62*).

## Data Availability and Code Availability

All data that support the findings of this study are available from the corresponding author on reasonable request.

All codes for the data analysis are available at GradinaruLab/SCN‐VIP‐estrous‐cycle: Anat Kahan, Jan 2023 (github.com)

## Acknowledgments

We would like to thank the Gradinaru lab for helpful discussions; Nikhila Swarna, Shinae Park, and Nathan Appling, for their technical assistance; Prof. Greg Anderson for the GnRH-Ab; Prof. Lance Kriegsfeld for the helpful discussions; and Dr. Catherine Oikonomou for the helpful discussions and manuscript review. This work was funded by the Heritage Medical Research Institute, the Vallee Foundation, and the Center for Molecular and Cellular Neuroscience in the Tianqiao and Chrissy Chen Institute for Neuroscience at Caltech (to V.G.). A.K. was supported by a Caltech Biology and Biological Engineering divisional postdoctoral fellowship and the Hebrew University Postdoctoral Fellowship for Women, Israel. A.K. acknowledges the support of her husband and children. A.K. and M.B were partially supported by the Global Grand Challenges grant from Bill & Melinda Gates Foundation. G.M.C. was supported by a PGS-D scholarship from the National Science and Engineering Research Council (NSERC) of Canada. B.W.B. acknowledges support from the Moore Distinguished Scholar Program at Caltech.

## Author contributions

A.K., G.M.C., M.B., and V.G. conceived the study. A.K., G.M.C., and M.B. performed the experiments, B.W.B. and A.K. developed the ML mathematical approach. A.K. analyzed and visualized the data. A.K. wrote the first draft of the manuscript. All authors read, reviewed, edited, and approved the final version of the manuscript.

