## Supplemental materials for "Dysregulated mammalian estrus cycle rescued by timed activation of VIP neurons in the circadian pacemaker and late afternoon light exposure"

### Supplementary materials

### Materials and Methods:

| REAGENT or RESOURCE | SOURCE | IDENTIFIER |
|---|---|---|
| **Antibodies** | | |
| GFP (Chicken); Cleared tissue: 1:200 | Aves | GFP-1020,<br>RRID AB_1608076 |
| GFP (Goat); brain slices: 1:1000 | abcam | ab6673<br>RRID AB_305643 |
| VIP (Rb); brain slices: 1:500 | Immunostar | 20077<br>RRID AB_572270 |
| c-Fos (Rb); Cleared tissue: 1:200; brain slices: 1:500 | abcam | ab209794<br>RRID AB_2905616 |
| VPAC2 (Rb) ; brain slices: 1:200 | Antibodies.com | A96020 (G790)<br>RRID AB_10695996 |
| KISS1R (Rb) ; brain slices: 1:40 ** | Proteintech | 15505-1-AP<br>RRID AB_10640578 |
| GnRH (GP) ; brain slices: 1:10000 | Greg Anderson, Otago University | GA-04 |
| Alexa Fluor 488 AffiniPure F(ab')2 Fragment Donkey Anti-Chicken IgY (IgG) (H+L)* | Jackson Immune | 703-546-155<br>RRID AB_2340376 |
| Alexa Fluor® 647 AffiniPure Fab Fragment Donkey Anti-Rabbit IgG (H+L)* | Jackson Immune | 711-607-003<br>RRID AB_2340626 |
| Cy™3 AffiniPure Donkey Anti-Guinea Pig IgG (H+L)* | Jackson Immune | 706-165-148<br>RRID AB_2340460 |
| Donkey Anti-Goat IgG H&L (Alexa Fluor® 488)* | abcam | ab150129<br>RRID AB_2687506 |
| **Bacterial and virus strains** | | |
| AAV5.hSyn.DIO.hM3D(Gq).mcherry | Addgene | 44361-AAV5 |
| AAV5.Flex.taCasp3.TEVp | UNC Vector Core | #45580 |
| AAV5.Ef1a.DIO.eYFP.WPRE.hGH | Addgene | 27056-AAV5 |
| AAVrt.CAG.FLEX.tdTomato.WPRE | Addgene | 51503-AAVrg |
| AAVrt.CAG.sgVipr2(A) | Glab | |
| AAVrt.CAG.sgVipr2(B) | Glab | |
| AAVrt.CAG.sgKiss1r | Glab | |
| **Chemicals, peptides, and recombinant proteins** | | |
| Normal Donkey Serum | Jackson Immune | 017-000-121 |
| Dimethyl sulfoxide | Fisher | D128-4 |
| Heparin | Sigma-aldrich | H3393-100KU |
| Triton X 100 | Sigma-aldrich | 93443-100ML |
| TWEEN 20 | Sigma-aldrich | 3005 |
| Glycine | Sigma-aldrich | G7126-100G |
| D-Fructose | Sigma-aldrich | F0127-1KG |
| α-Thioglycerol | Sigma-aldrich | M1753-100ML |
| Clozapine N-oxide (CNO) dihydrochloride | Tocris | 6329/10 |
| **Recombinant DNA and virus perpetration components** | | |
| pSpCas9(BB)-2A-GFP, Feng Zhang | Addgene | 48138 |

| | | |
|---|---|---|
| AAV2-retro, Alla Karpova and David Schaffer | Addgene | 81070 |
| Neuro-2a cells | ATCC | CCL-131 |
| HEK293T cells | ATCC | CRL 3216 |
| QuickExtract DNA Extraction Solution | Lucigen | QE09050 |
| NEBuilder HiFi | New England Biolabs | E2621 |
| polyethylenimine | Polysciences | 24765-1 |
| Iodixanol, Optiprep | Sigma-Millipore | D1556 |
| **Software and algorithms** | | |
| ICE analysis | Synthego | |
| ImageJ, version 2.0.0-rc-69/1.52n | NIH, open source | https://fiji.sc/ |
| Adobe Illustrator, 24.0.2 (64-bit) | Adobe | https://www.adobe.com/products/illustrator.html |
| Matlab R2020b, R2018a | MathWorks Inc. | https://www.mathworks.com/products/matlab.html |
| Imaris 9.2.0 | Oxford Instruments | https://imaris.oxinst.com/ |
| Zen (LSM 880, 2.3 lite) | Zeiss Microscopy | https://www.zeiss.com |
| BioRender | BioRender | https://biorender.com/ |
| Custom code to analyze and generate figures | This paper | GradinaruLab/SCN-VIP-estrous-cycle: Anat Kahan, Jan 2023 (github.com) |
| **Other** | | |
| Optical fiber, 400 µm diameter, 7 mm long | Doric Lenses | MFC_400/430-0.48_7mm_ZF1.25_FLT |
| Mono Fiberoptic Patch cable (fiber photometry) | Doric Lenses | MFC_400/430_0.48_2m_FC_ZF1.25_FL |
| Double patch cable (optogenetics) | Doric Lenses | BFP(2)_400/430/1100_0.48_1.5m_FCM*-2xZF1.25(F) |

Table S1: Detailed reagent and resources.

* All secondaries were used at 1:200 for cleared tissue, and 1:1000 for 70/100µm brain slices

**This ab was incubated for two over-nights in 4°c.

| sgRNA | crRNA sequence | Forward primer | Reverse primer |
|---|---|---|---|
| Vipr2-B | GGGCATCCGAATGACCCACC | TTGACTCACCCAGGAAAGCC | TTGGAAATGGGCAGCAGACT |
| Kiss1r-B | AGCGTTGGACGGAGCCCACC | CAGAGGAGCCTCTTTCCAGC | TTCCTGACTTGGACCCCAGA |

Table S2: related to figure 4. crRNA sequences and associated primers for Genetic knockout in GnRH neurons using CRISPR

##### *Experimental animals*

Mice used in this work include C57BL/6NCrl (Charles River), VIP-IRES-Cre (Jackson Laboratory Stock, JAX, 010908) crossed to C57BL/6NCrl (Charles River), or Ai162 (GCaMP6s reporter line, JAX, 031562). The usage of Ai162 line, when crossed to a Cre line, allows suppression of GCaMP6s expression using a doxycycline diet. Therefore, for fiber photometry recording, the diet was given to the breeding pair and the offspring until 2-4 weeks before surgeries, to prevent toxicity or developmental issues. For optogenetic experiments, VIP-IRES-Cre line was crossed to Ai132 (ChR2, JAX, 24109). For GnRH neurons gene editing, GnRH-Cre line (Gnrh1-Cre, JAX, 21207) was crossed to Rosa26-Cas9 (JAX, 24858). Animals were group housed (2-4 per group) whenever possible to ensure a regular estrous cycle. Mice were single-housed for FP experiments that required 24/7 recordings and optogenetic experiments. Females that were single housed for calcium recording received a handful of fresh male mice bedding every cage change to ensure regular estrous cycles. Mice had a running wheel or a cleared tube for enrichment in their home cage. For optogenetics, mice were connected with two-branched patch cables, so running wheels/tubes were impossible. Mice were kept at 12:12 light cycle, unless stated otherwise. The ambient room light intensity was 4.49E14 photons/cm$^2$/sec (150 lux). Light at 'lux' units was measured with the Light Meter smartphone app ('My mobile Tools Dev'). Light power (mW) was measured with Thorlabs S120C photodiode. The energy units were divided by the sensor dimensions, 0.94 cm$^2$. Unit conversion from mW/cm$^2$ to photons/cm$^2$/sec were calculated using: $n_{photons} \cdot E = n_{photons} \cdot \frac{hc}{\lambda}$, where $h$ is Plank constant, c is the speed of light, and $\lambda$ is the light wavelength (at maximum). Mice were used until 10 months old to ensure a regular estrous cycle (*1*), and were age-matched for each experiment. Animals had *ad libitum* access to food and water.

FP recording, light manipulation, optogenetic and chemogenetic experiments were performed in rat cages (40 x 34 cm). For FP, the lid was modified to have a hole in the middle for the patch cable.

Light manipulation experiments with PIR detection and/or optogenetic manipulation were performed in rat cages, with a 10-inch diameter Plexiglas cylinder, 34 cm in height, to ensure sufficient PIR detection. Mice were group housed for light manipulation experiments. For optogenetic experiments, mice were single-housed due to the patch cable. Other light manipulation experiments (cohort 1 of DREADD, C57) were performed in regular mice cages.

Animal husbandry and experimental procedures involving animal subjects were conducted in compliance with the Guide for the Care and Use of Laboratory Animals of the National Institutes of Health and approved by the Institutional Animal Care and Use Committee (IACUC) and by the Office of Laboratory Animal Resources at California Institute of Technology under IACUC protocol 1739. Mice were excluded from the entire FP experiment if there was no dynamic photometry signal or no two-photon signal 3 or 5 weeks after surgery, respectively.

***Surgery***

*Stereotactic viral vector injections* were made in mice anesthetized with isoflurane (5% induction, 1–1.5% maintenance) and placed on a stereotaxic frame (942, David Kopf Instruments, CA, USA). An incision was made to expose the skull, including Bregma, lambda, and the target sites' external coordinates. Stereotaxic coordinates were measured from Bregma and were based on the Mouse Brain Atlas (*2, 3*), and improvement was made based on 2D or LiGS histology. A craniotomy hole was drilled above the target. Virus injection and implantations were performed as follows:

*Cell apoptosis:* VIP-Cre mice were injected with AAV5.Flex.taCasp3.TEVP virus (Experimental, 2.9E12 VG/ml) or AAV5.Ef1a.DIO.eYFP.WPRE.hGH (control, 2.3E12 VG/ml), AP -0.3/-0.2 mm, ML ± 1.19 mm, DV -5.75 and -5.6 mm from the brain, left and right sides, at 13 degrees, 2 x 350 nl each side.

*Excitatory DREADD*: VIP-Cre mice were injected with excitatory DREADD or control virus at the SCN; AAV5.hSyn.DIO.hM3D(Gq).mcherry (1.0E13 VG/ml) or AAV5.Ef1a.DIO.eYFP.WPRE.hGH (1.0E13 VG/ml) respectively, AP -0.3/-0.2 mm, ML ± 1.19 mm, DV -5.75 and -5.6 mm from the brain, left and right sides, at 13 degrees, 2 x 300 nl each side.

*Genetic ablation in GnRH neurons:* GnRH-Cre x Cas9 mice were injected with AAVrt.U6-sgRNA(SapI)-CAG-mRuby2-WPRE-hGHpA (1.4E13 VG/ml) or AAVrt.CAG.flex.tdTomato (6.4E12 VG/ml, control), into the median eminence, targeting GnRH processes (*4*), prepared as described in the following section. AP -1.4 mm, ML ± 0.3 mm, DV -5.65 and -5.4 mm from the brain, left and right sides, at 0 degrees, 2 x 300 nl each side.

All viruses were injected at a rate of ~80 nl/min using a blunt 33-gauge microinjection needle within a 10 µl microsyringe (NanoFil, World Precision Instruments, WPI) by an UltraMicroPump (UMP3-4, WPI), controlled by a pump controller (Micro4, WPI)

*Fiber photometry (FP), Optogenetics*: Two optical fibers with a cut length of 7 mm and diameter of 400 µm (MFC_400/430-0.48_7mm_ZF1.25_FLT ,NA=0.48, Doric lenses, Canada) were firmly mounted to a stereotaxic holder. Two fibers were implanted to improve the probability of successfully targeting the structure. A thin layer of Metabond (Parkel) was applied to the skull surface to secure the fiber. In addition, a thick layer of black dental cement (JET denture repair powder and liquid) was applied to secure the fiber implant and to prevent interferences between the excitation light to light-sensitive targets that are not the SCN.

Extended details about viruses and implants can be found in Table S1.

In some cases, following implant surgery, an ovariectomy (OVX) was performed as follows:

*OVX:* Each mouse was given a single dose of ketoprofen 5 mg/kg SC and sustained-release buprenorphine at 1mg/kg SC. The mouse was then anesthetized with 1-5% isoflurane in an induction box followed by maintenance in a nose cone and was maintained on a heating pad throughout the surgery. A small dorsal midline incision was made over the abdomen. The abdominal cavity was entered via a blunt puncture through the abdominal wall. The ovary was dissected. The fat pad and tissue were returned to the abdominal cavity, and the abdominal wall was closed with 4-0 absorbable multifilament suture in an interrupted pattern. The process was repeated on the opposite side through a single incision. The skin incision was closed with surgical wound clips or by suturing with a monofilament 4-0 suture material in an interrupted pattern. Before closure, bupivacaine (1 mg/kg of 0.25% solution) was applied subcutaneously to the wound margins. Mice received 30mg/kg Ibuprofen *ad lib* (20 mg per 100 ml water) for at least 5 days. For all OVX females, surgery success was verified by collecting vaginal smears for at least 10 days, showing either diestrus or metestrus states.

All mice were given 1 mg/kg sustained-release buprenorphine and 5 mg/kg ketoprofen s.c. Intraoperatively and received 30 mg/kg ibuprofen p.o. in their home cage water for at least five days postoperatively for pain. Mice were allowed a minimum of 14 days for surgical recovery before participation in behavioral studies.

***Genetic knockout in GnRH neurons using CRISPR***

*Guide RNA cloning and validation:* sgRNA sequences were validated using pSpCas9(BB)-2A-GFP (PX458; a gift from Feng Zhang, Addgene ID: 48138). Briefly, Neuro-2a cells (ATCC CCL-131) were transfected in duplicate with PX458 containing sgRNAs of interest, and their DNA was extracted at 72 hours post-transfection (QuickExtract DNA Extraction Solution, Lucigen, QE09050). The targeted region was amplified with PCR, and Sanger sequenced. Reads were analyzed for editing efficiency using ICE analysis (Synthego). Only sgRNA sequences that yielded >70% editing efficiency (roughly corresponding to the transfection efficiency) were used for *in vivo* experiments. Extended data regarding Recombinant DNA and virus perpetration components information can be found in Table S1. crRNA sequences and associated primers can be found in Table S2.

For *in vivo* sgRNA delivery, pAAV.U6-sgRNA(SapI)-CAG-mRuby2-WPRE-hGHpA was generated by inserting a dsDNA fragment containing the U6 promoter and *Streptococcus pyogenes* Cas9 guide RNA scaffold with SapI sites for crRNA insertion into MluI-digested pAAV-CAG-mRuby2 (*5*), using NEBuilder HiFi (New England Biolabs, E2621). crRNA sequences were inserted by annealing complementary oligos containing overhangs, followed by ligation into SapI-digested pAAV.U6-sgRNA(SapI)-CAG-mRuby2-WPRE-hGHpA.

*AAV production:* pAAV.U6-sgRNA-CAG-mRuby2-WPRE-hGHpA containing validated sgRNA sequences was packaged into AAV2-retro (*6*) capsids through triple transient transfection of HEK293T cells (ATCC, CRL 3216) with polyethylenimine (Polysciences, 24765-1), followed by cell lysis and purification over iodixanol (Optiprep; Sigma-Millipore, D1556), as previously described (*7*). The AAV2-retro rep-cap plasmid was a gift from Alla Karpova and David Schaffer (Addgene ID: 81070). Viral titers were obtained through qPCR, using a linearized genome plasmid as a standard. Cells were verified to be free of Mycoplasma contamination prior to AAV production.

##### *Fiber photometry recording*

FP is a method for measuring population calcium-dependent fluorescence from genetically-defined cell types in deep brain structures using a single optical fiber for both excitation and emission in freely moving mice. A detailed description of the system can be found elsewhere (*8*). Briefly, our system employed a 490 nm LED for fluorophore excitation (M490F1, Thorlabs; filtered with FF02-472/30-25, Semrock) and a 405 nm LED for isosbestic excitation (M405F1, Thorlabs; filtered with FF01-400/40-25, Semrock), which were modulated at 211 Hz and 531 Hz, respectively. Two systems were used for recording, both controlled by a real-time processor (System 1: RX8-2; System 2: RZ5P, Tucker-Davis Technologies), and delivered to the implanted optical fiber via a 0.48 NA, 400 µm diameter mono-fiber optic patch cable (MFP_400/430/LWMJ-0.48_2 m_FC-ZF1.25, Doric Lenses). The emission signal from isosbestic excitation, which was previously shown to be calcium-independent for GCaMP sensors (*9, 10*), was used as a reference signal to account for motion artifacts and photobleaching. Emitted light was collected via the patch cable, collimated, filtered after passing through a focusing lens (System 1: MF525-39 filter, Thorlabs, 62–561 focusing lens, Edmunds Optics; System 2: Mini Cube FMC6, Doric Lenses), and detected by a femtowatt photoreceiver (Model 2151, Newport). Photoreceiver signals were demodulated into GCaMP and control (isosbestic) signals, digitized (sampling rates: System 1: 382 Hz; System 2: 6 Hz), and low-pass filtered at 25 Hz using a second-order Butterworth filter with zero phase distortion. A least-squares linear fit was applied to align the 405 nm signal with the 490 nm signal. Then, the fitted 405 nm signal was subtracted from the 490 nm channel and then divided by the fitted 405 nm signal to calculate dF/F values.

##### *Behavioral assays*

*Estrous Cycle Stage Identification:* Estrous cycle stage was recorded following established protocols (*11*). Briefly, vaginal smears (VS) were collected with a 100 or 20 µl pipette holding 15 µl saline at ZT10-14. In cases where the room was entered for injections or light stimulation, VSs were taken at the time of interference to reduce entrainments. VS were flattened and allowed to dry on a microscope slide (Adhesion Superfrost Plus Glass Slides, Brain Research Laboratories) and stained with cresyl violet dye (0.1%, Sigma-Aldrich, C5042-10G). Smears were then analyzed using light microscopy. Proestrus is characterized by a high number of nucleated epithelial cells. The estrus stage has a high number of cornified epithelial cells. During metestrus there is an increased amount of leukocytes, with the presence of cornified epithelial cells and some nucleated epithelial cells. Diestrus is characterized by mostly leukocytes, with low presence of cornified epithelial cells and nucleated epithelial cells. We defined the number of proestrus events based on the number of identified proestrous events, even if a full cycle was not detected during the monitored period.

*OVX and hormonal replacement:* OVX was verified by having a VS profile of diestrus or metestrus. At least four weeks after OVX, sex hormone stimulation was performed as follows; estradiol was administered at 10 μg and progesterone at 500 μg in 0.05 ml of sesame oil delivery s.c. at ZT8, which has been shown to induce sexual receptivity. These estradiol levels are comparable to physiological proestrus peak levels (*12*).

*Locomotor activity (LMA) detection:* Mice were placed in a rat cage (40 X 34 cm). A PIR motion detector (COMPASS (*13*)) was placed 35 cm above the cage bottom, and a Plexiglas cylinder was placed in the middle (10-inch diameter) to ensure even detection of mouse activity. Mice were placed with a tube used for handling to reduce anxiety (*14*).

*Chemo- and Optogenetics:* After recovery from the surgery, mice were moved to the behavioral room with the light cycle used for the experiment and handled for at least two weeks before the experiment started. Female mice were placed in a rat cage with a round cylinder, as described above. Mice were given at least 3-4 days to acclimate to the new cage setup and handling. Following the first baseline session, the light cycle was changed to DD, with additional light stimulation at CT0 (DD +$CT0_{0.5L}$), to preserve their locomotor rhythm as much as possible. At either CT4 or CT10, stimulation was given as follows:

For optogenetics: a 447nm laser (MDL-III-447-200mW, OptoEngine LLC) was used, at 15 Hz, pulse duration of 15 ms, and an intensity of 10 mW, repeated for one-hour total. This pattern was adopted from Mazuski et al. (high frequencies pattern) (*15*). In addition to the black dental cement, we used two layers of black heat-shrink tubes to prevent interferences between the relatively high-intensity laser and light-sensitive targets, which are not the SCN.

For DREADD: CNO was prepared in distilled water at 1 mg/ml as a stock solution and kept for up to a month at -20 °C. CNO was injected i.p. at 0.1 mg/ml, (0.1 ml per 10 gr) for 1 mg/kg.

To validate opto- and chemogenetic activation *post hoc*, we activated the cells for one hour under the same conditions or injected CNO one hour before mice were sacrificed (at ZT14) and examined c-Fos and ChR2 or hM3Dq expression using IHC.

During all light and neuronal manipulation studies, vaginal smears (VSs) were taken daily at ~ZT10, except for opto- and chemogenetic experiments, in which VSs were taken prior to stimulation to prevent further interference. For these experiments, females were assessed for three weeks for a regular estrous cycle using vaginal smears, and their weight was monitored every three weeks. Females with low number of cycles, <3 over three weeks, were removed from the cohort, but left in the cage for social stability. In one case, a female was excluded from the experiment due to severe weight loss (>10%). For light and $SCN^{VIP}$ activation experiments, females that were not affected by the reduced light conditions were not considered as 'rescuable', and therefore were excluded (6/41).

*Released egg collection*: Ovulating eggs were collected from the oviduct the day after ovulation, within 24-30h after the detected proestrus state. Cumulus-oocyte complexes were isolated in M2 media (Sigma, M7167) supplemented with 0.3% hyaluronidase (Sigma, H3506) to determine the number of eggs ovulated.

##### *LiGS 3D histology*

*LiGS tissue preparation:* LiGS histology of implanted mice was performed as described previously (*16*): after perfusion (20 ml 1× PBS followed by 20 ml 4% PFA), the implant was kept intact; the skin and the lower jaw were gently removed. The remaining skull, including both brain and implant, was placed in 4% PFA for two days. Following fixation, samples were washed in 1 x PBS, then placed in 15% and 30% sucrose solution for cryoprotection. For LED coupling, samples were placed in 22x22 mm disposable embedding molds (70182, EMS) with OCT (Tissue-Tek Compound, Sakura Finetek) and were frozen with an ethanol/dry-ice bath (−78°C). The brain was positioned such that the optical implant was perpendicular to the cube during freezing. Next, a 5 mm LED (Chanzon, yellow) was placed directly above the optical device and secured with additional OCT. To give the brain–OCT cube a flat surface, a 20x40 mm embedding mold (70184, EMS) was filled with OCT while the brain (and coupled LED) was placed upside down and dipped together into the ethanol/dry-ice bath. This process created a large, stable OCT cube that included the sample and the coupled LED. The sample was then cut on one side to expose the LED wires and stored at −80°C until needed.

*Light-guided cryo-sectioning:* Brains in OCT were placed in a cryostat and sliced from the bottom in 50–100um steps with the LED turned on. To ensure a reproducible light intensity, we used a power supply (DG1022, RIGOL) set to 2.1 V. We used the profile of the light spread by manual observation to define the sectioning endpoint: after the scattered light became sharp, sectioning was continued in small steps (20–50 μm) until a shaded area was seen in the fiber location when the LED was turned off. After sectioning, samples were left at room temperature (RT), allowing the OCT to melt gradually. Samples were then gently placed in a tube and washed with 1× PBS solution.

*LiGS staining with IHC:* After sectioning, brains were put in 4% PFA for 1–3 hours for additional fixation. The staining protocol was adapted from the iDisco protocol without the pretreatment step [20]. Briefly, the samples were incubated for two days at 37°C in permeabilization solution, followed by two days in blocking solution at 37°C. Next, the samples were incubated with primary antibodies at a 1:200 concentrations for 5–7 days in PTwH/5%DMSO/3%donkey serum, at 37°C. After washing at room temperature (RT) until the next day, samples were incubated with a secondary antibody in PTwH/3% donkey serum at 37°C for 5–7 days. Lastly, samples were washed with PTwH at RT until the next day.

*LiGS Clearing*: Samples were cleared using the SeeDB protocol at RT (*17*).

##### *Thin slice staining:*

After perfusion (20 ml 1× PBS followed by 20 ml 4% PFA), brain sections (70 or 100 μm, Leica Vibratome VT1200S) were first incubated in a blocking buffer solution (1× PBS solution with 0.1% Triton X-100 and 10% normal donkey serum) for at least one hour, followed by incubation with primary antibodies in the blocking buffer solution at 4°C overnight. The next day, sections were thoroughly washed four times in 1× PBS (15 min each). Next, the brain sections were transferred to a blocking buffer solution with secondary antibodies and left overnight at 4°C or for two hours at RT. Next, sections were washed as described above and mounted on glass microscope slides (Adhesion Superfrost Plus Glass Slides, Brain Research Laboratories). After the sections were completely dry, they were cover-slipped after applying mounting media (VECTASHIELD® PLUS).

Extended information about IHC materials can be found in Table S1.

##### *Histological imaging*

Histological images were obtained with Zeiss 880 confocal microscope at 10X and 40X objectives. Images were analyzed in ImageJ, Matlab, and/or Imaris (Bitplane). Cell identification of GnRH neurons in the MPA was performed with the "Spots" function in Imaris (9.8.0 and 9.9.1), with a diameter of 15 μm, for each channel separately. In addition, the "Statistics" toolbox of Imaris was used to identify overlapped populations (<10 μm). For each animal, three brain slices which included the MPA were quantified, and the injection quality was validated by observing the injection site (Figure S6).

##### *Data analysis*

*Light detection*

*LMA:* Following PIR motion detection, the mean activity and onset were calculated using an angular presentation of each 24-hour period, using the "CircStat" toolbox for circular statistics (Matlab (*18*)). Slopes were calculated using a linear regression function, looking at the mean activity or onset values over days (Matlab, with 95% confidence).

*Fiber photometry (FT):* dF/F was first aligned using the 405 nm channel, as previously used [30]. When z-scored data was presented, we used: $dF/F = F - median\,(F_b)/mad\,(F)$ (where 'mad' is the median absolute deviation, and $F_b$ is the background signal during the dark phase). The event rate was calculated using the peak finder algorithm ("findpeaks" using "Annotate," Matlab), with a threshold of 1.2 std above the mean value at the baseline period for the 10 minutes/h sessions, and 0.4 for the ZT10-13 recording, justified by the reduced ability of 'event rate' analysis to identify low amplitude frequencies, shown by the FFT classification analysis. For 10 minutes/h sessions, data was collected starting at ZT8 for 24h. For the analysis, data were shifted by -8 hours. That way, the estrous cycle definition was accurate for the last 8 hours of the light phase, followed by the rest of the light phase and the following dark phase. In the case of proestrus, ovulation occurs during the dark phase, and the following light phase should be considered as estrus. For male-to-female comparison, female data was taken for the proestrus day.

*FT FFT analysis*: For each dF/F signal (10 minutes/h), FFT was calculated ('fft', Matlab). For each hour, integrated FFT power values were calculated for one set of frequency intervals: [0.0033 0.007; 0.007 0.05; 0.05 0.1; 0.1 0.25; 0.25 0.45; 0.45 1.0; 1.0 1.35] Hz. The lower limit was set to 0.0005 for the ZT10-13 recording, due to the longer time frame. FFT autocorrelations were calculated for each FFT power interval over the coarse frequency intervals (set (a),'autocorr', Matlab). See Figure S3 for an example of the FFT approach on synthesized data.

For classification, a spectrogram of the fine frequency intervals was prepared for each session. Each session was tagged by sex and by hormonal state. Prior to machine learning, the spectrogram dimensions were reduced by (1) choosing only frequency ranges between 0.003 to 1 Hz (the first six intervals), (2) applying PCA (pca, 'Matlab'), i.e. 24 h x 6 frequencies intervals were reduced from 144 to 12, (3) when indicated, only a subset of the recording 10-minute intervals were used. A leave-one-out cross-validation (LOOCV) approach was used: in each step, one session was left out to be the 'testing-set', while the rest of the sessions were used as a 'training-set', after randomly choosing the same number of sessions for each state. For training, we used either a "support vector machine" (SVM, 'fitcecoc', Matlab), or a "discriminant analysis" ('fitcdiscr', Matlab) classifier, followed by prediction ('predict' Matlab). In each classification, the prediction rate was calculated for the ability to predict each of the two states, averaged over 5 (24h dataset) or 10 cycles (ZT10-13 dataset), and the score was plotted as a matrix, weighted by color ('matvisual', Matlab, created by Hristo Zhivomirov). SEM between cycles was 1% on average.

Extended information about Software and algorithms can be found in Table S1.

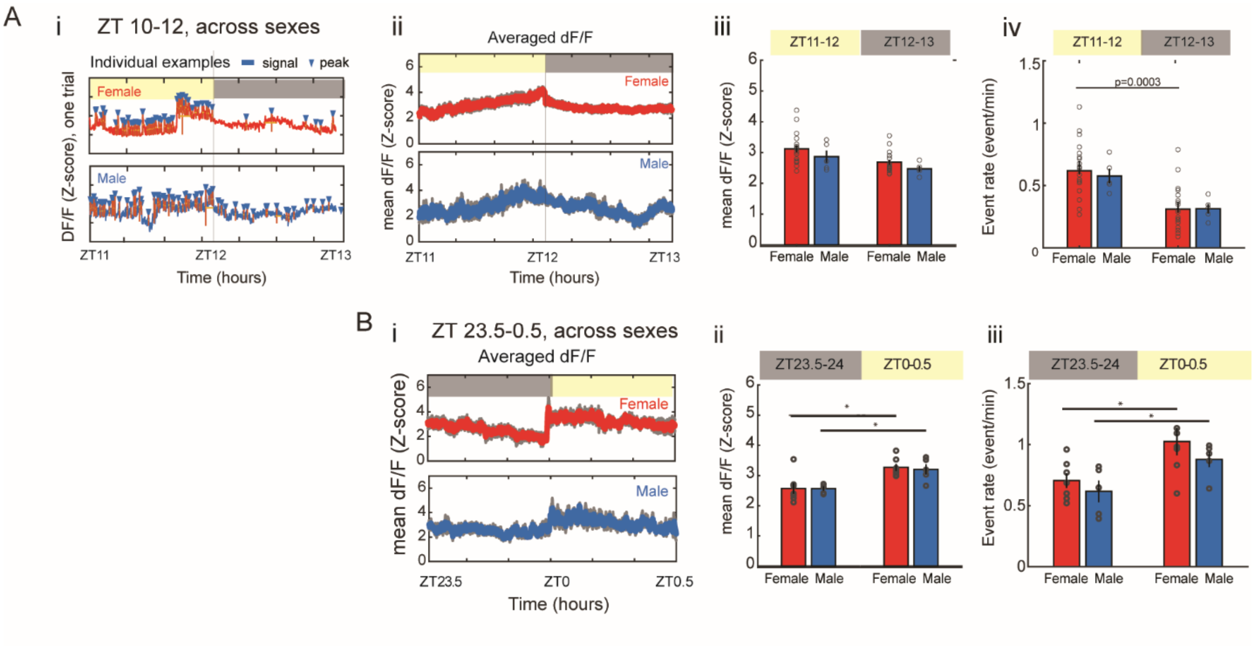


**Figure S1, related to Figure 2**: **FP activity of SCN$^{VIP}$ neuron at the transition from light to dark (ZT11-13) showing sex differences at event rates, but not at the transition from dark to light (ZT23.5 to 0.5).** (A) SCN$^{VIP}$ FP activity at ZT11-13 (i) Representative examples of FP recordings from a female (red) and male (blue) with peak identification (blue triangles). (ii) Averaged responses (mean±sem, sem in gray. females: n=14, males: n=5, averaged for at least three repeats each). (iii-iv) Mean dF/F and event rates, compared between males and females at ZT 11-12 and ZT 12-13 (mean±sem, with individual values shown). Event rates were higher during the light phase from ZT 11-12 compared to ZT 12-13 in females but not in males (0.58±0.06 and 0.64±0.05 events/min at ZT11-12 vs. 0.31±0.04 and 0.32±0.04 at ZT12-13, There was no significant difference in mean dF/F (3.1±0.1 and 2.69±0.08 a.u. at ZT11-12 vs. 2.9±0.2 and 2.47±0.09 a.u. at ZT12-13, males and females, respectively. (B) SCN$^{VIP}$ FP activity at ZT23.5-0.5. (i) Averaged responses (mean±sem, sem in gray, females: n = 7, males: n = 5, averaged for at least three repeats each). (ii) Mean dF/F, showing no differences between males and females and a significant increase from dark to light. (iii) Event rates show no differences between males and females and significantly increase from dark to light. Nonparametric Kruskal-Wallis test, * $p < 0.05$.

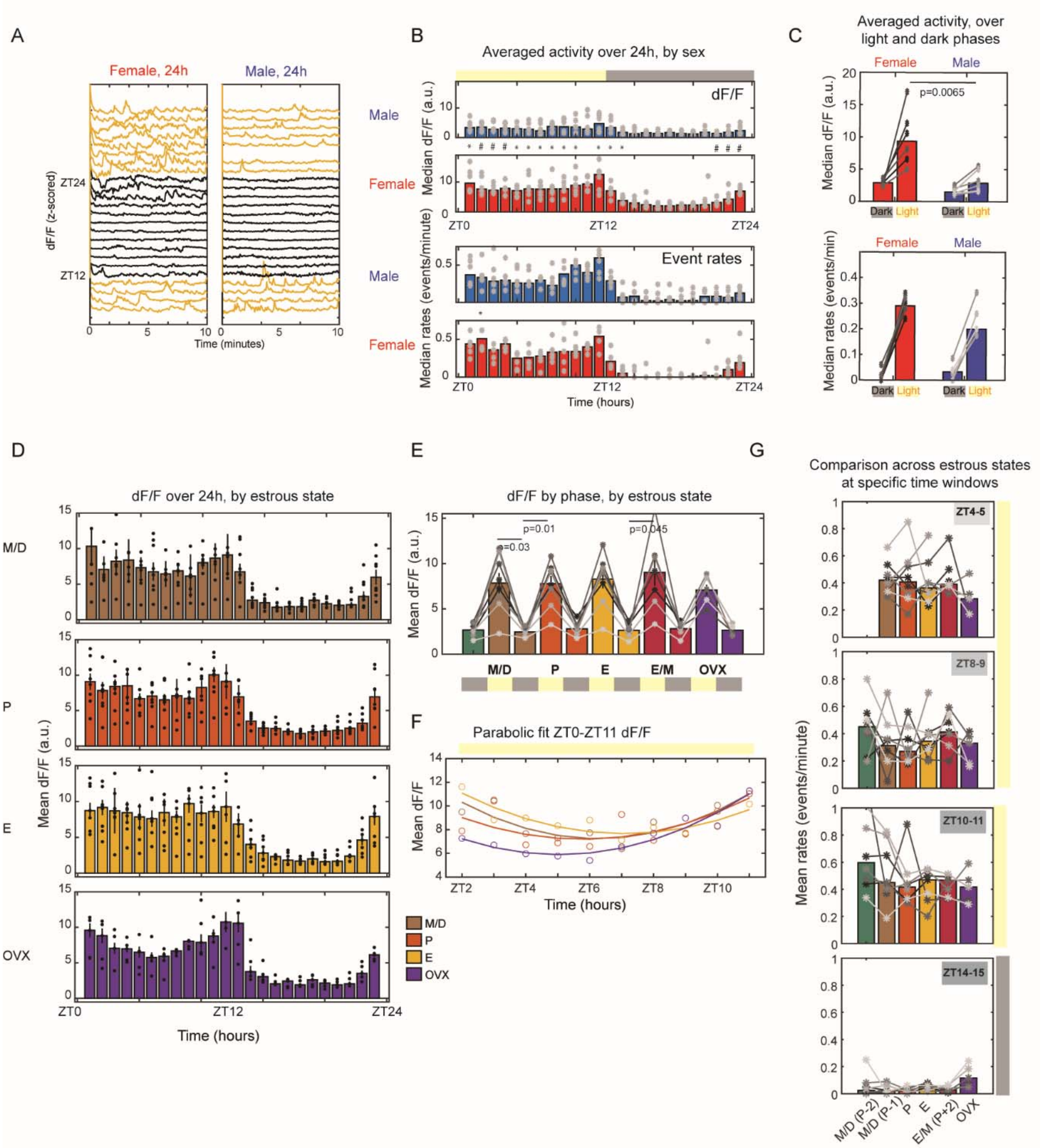


**Figure S2, related to Figure 2: Circadian changes in SCN$^{VIP}$ signal are sex-dependent but not estrous-cycle dependent.** (A) Representative examples of 10-minute-per-hour recording, over 24 hours, from a female (left) and a male (right) mouse. (B) DF/F (top) and event rates (bottom) over 24h of recording (p values compare females, n=6 (red) vs. males, n=6 (blue), at least 3 repeats each. (C) Comparison of median dF/F (top) and median event rate (bottom) between males and females over the dark and light phases. (D) DF/F over 24h of recording across estrous states (n=8, each state is represented at least three times in each female). (E) DF/F averaged over dark and light phases. For clarity, significance is marked only between adjacent periods. (F) Parabolic fits to median dF/F along ZT2-11. (G) Averaged event rates over two hours at different points during the day, showing no significant differences between estrous states. Data was assigned relative to the proestrus day, therefor M/D states were identified either one or two days before proestrus (M/D P-2 and M/D P-1), as well as two days after proestrus, which were either E or M. Nonparametric Kruskal-Wallis test, followed by Tukey's correction when multi-comparison was done (*p<0.05; **<0.01; # < 0.005).

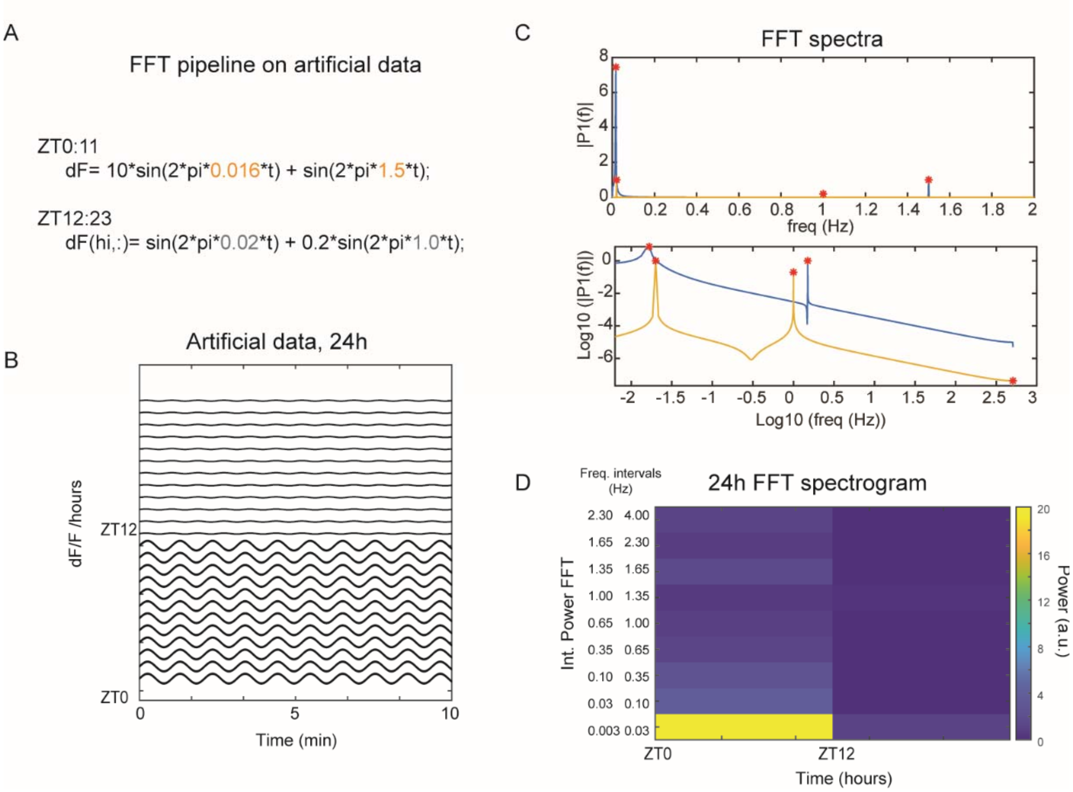


**Figure S3, related to Figure 2: FFT spectrogram creation approach using artificial data.** (A) Artificial FP data is a combination of two sine signals, for ZT0-11 and ZT12-23 separately, at frequencies of 0.016 and 1.5Hz, with amplitudes of 10 and 1, as well as 0.02 and 1.0 Hz, with amplitudes of 1 and 0.2, respectively. (B) FP artificial oscillations. (C) Power spectra of the artificial FT data ('fft', Matlab), using regular (top) and logarithmic (bottom) scales. Red dots indicate the contributing frequencies, 0.016, 0.02, 1.0, and 1.5 Hz. (D) Integrated FFT over different frequency intervals.

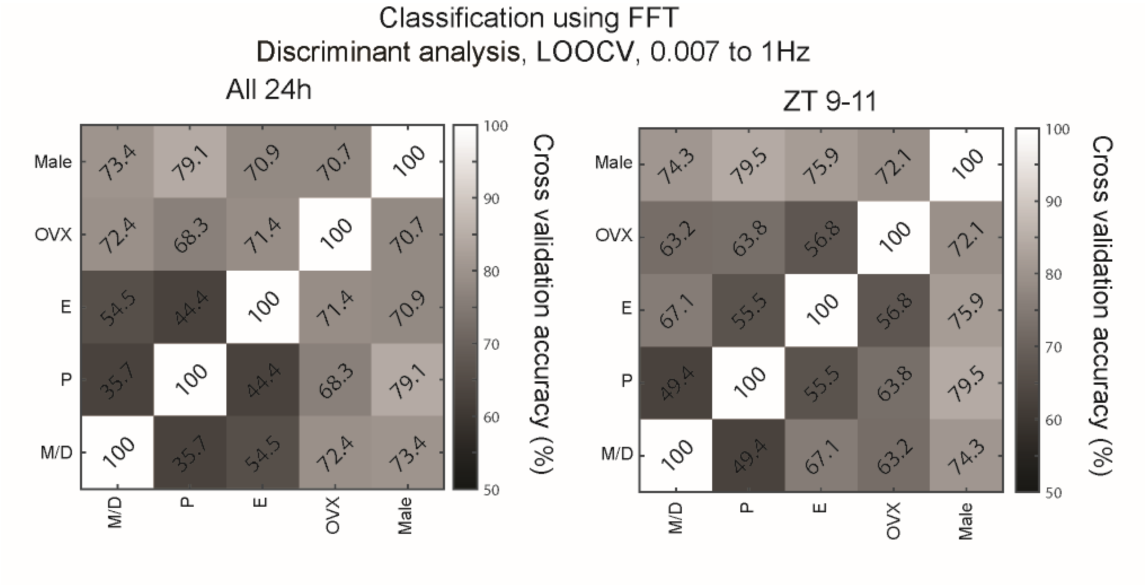


**Figure S4, related to Figure 3: Cross-validated accuracy with "Discriminate analysis" algorithm,** using the full 24h FFT spectrograms (left) or ZT9-11 (right). The accuracy values show similar prediction abilities to the SVM algorithm.

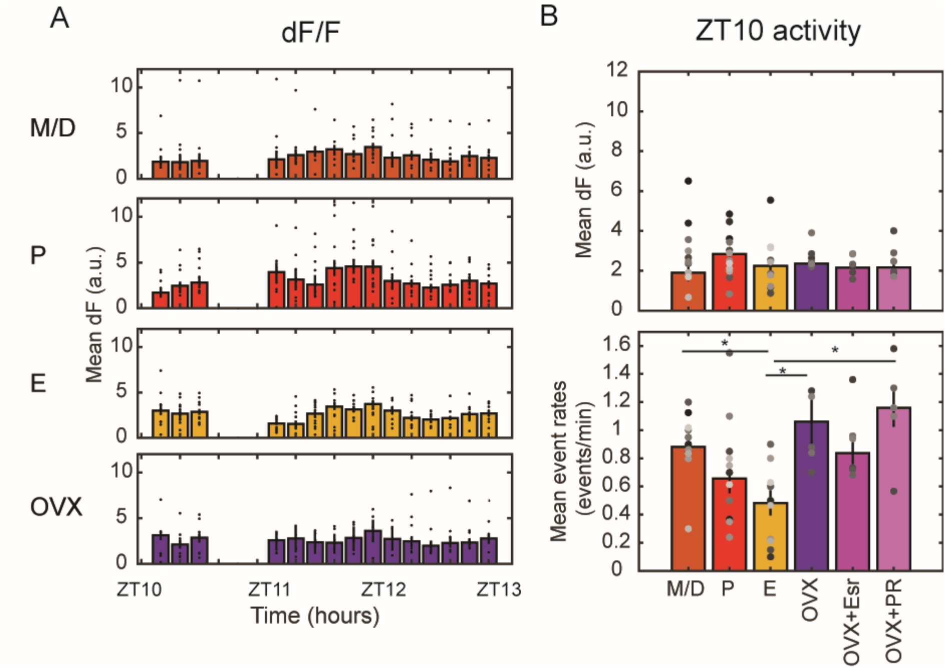


**Figure S5, related to Figure 3. Activity analysis of ZT10-13 FP data, including OVX induction with sex hormones**. (A) Averaged dF/F at ZT10-13 across estrous states. (B) Quantified dF/F (top) and event rates (bottom), including OVX induction with sex hormones. *$p<0.05$; Kruskal–Wallis test, Tukey's correction for multi-comparisons.

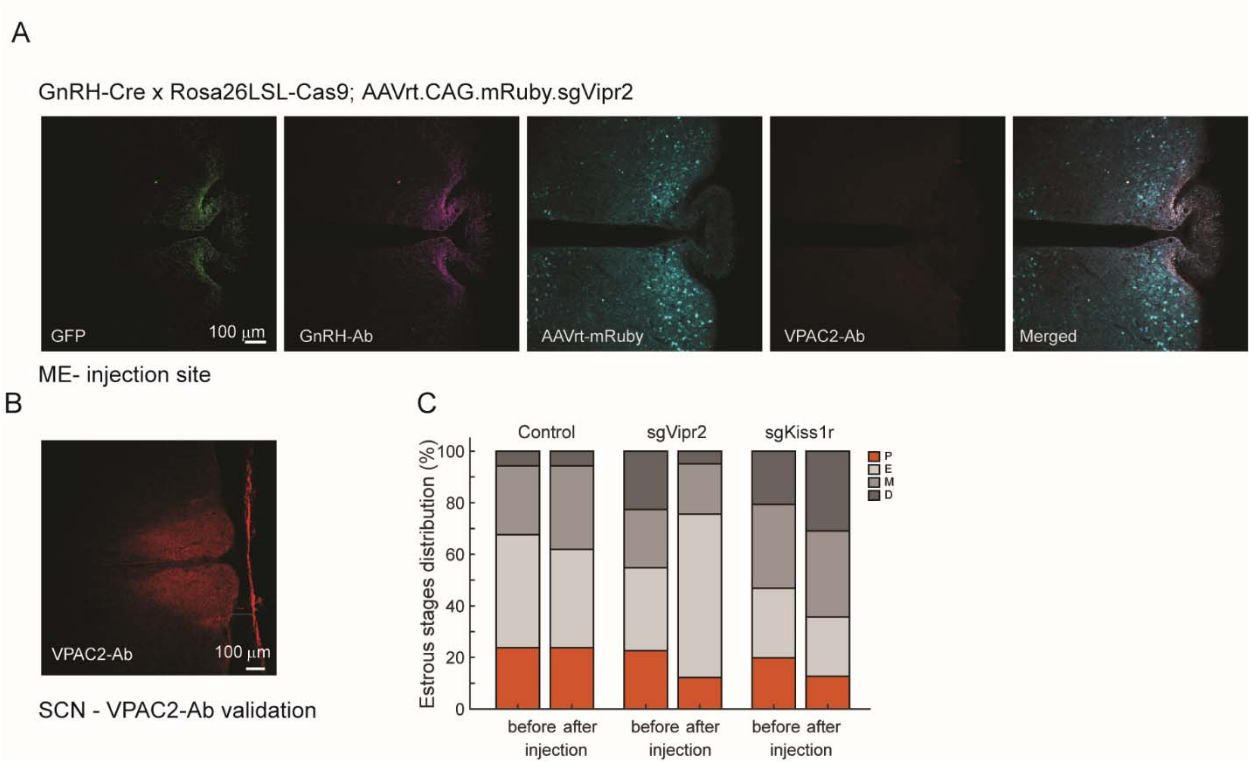


**Figure S6, related to Figure 4. Validation of GnRH-Cre x cas9 injection site and VPAC2 antibody and estrous state distributions**. (A) The injection site, the median eminence (ME). (B) VPAC2 Ab validation based on expression in the SCN. (C) Estrous states distributions over three weeks, before and after injection, control (Ctrl, n=5), sgVipr2 (n=4), and sgKiss1r (Exp, n=6).

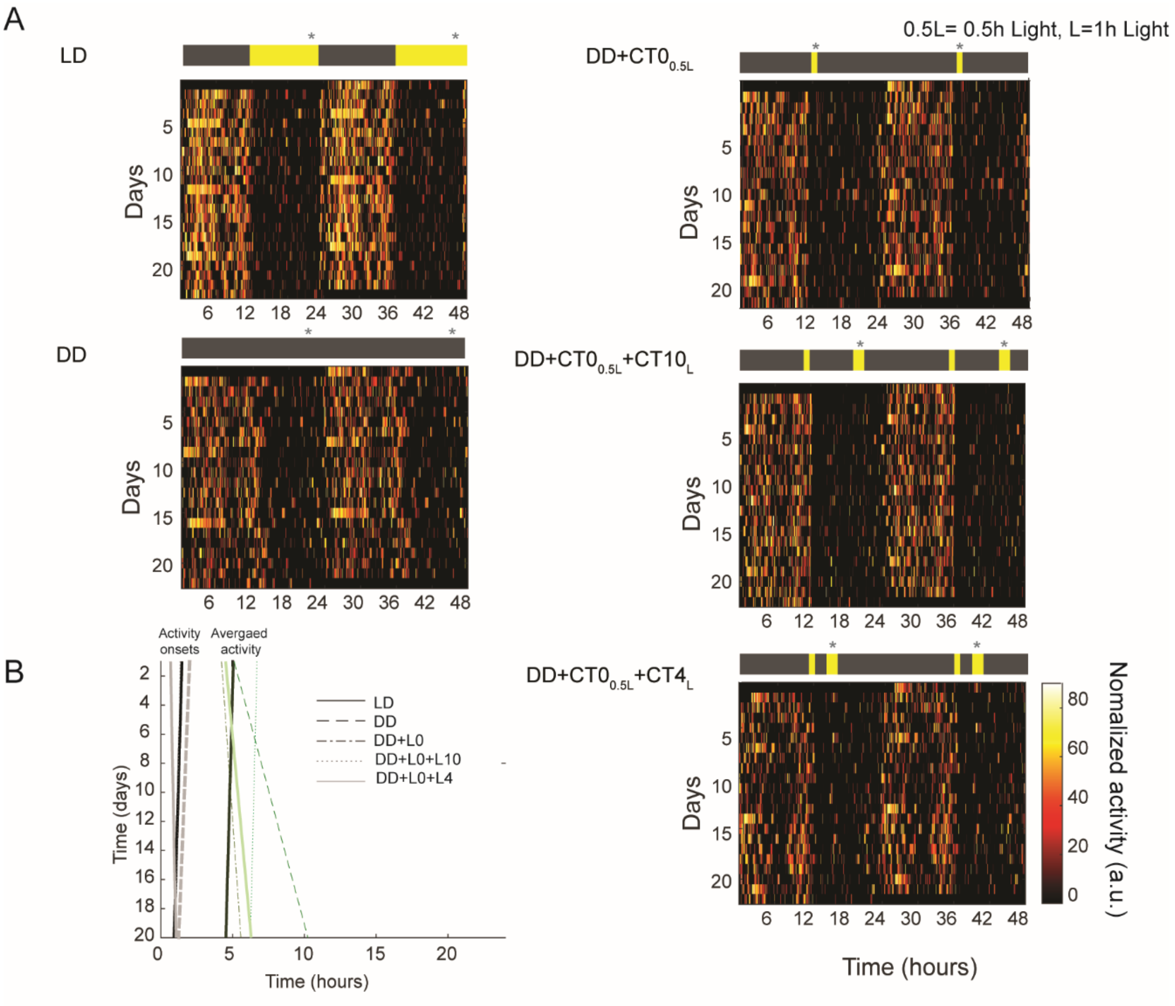


**Figure S7, related to Figure 5**. **LMA under different light conditions of VIP-cre female mice**. (A) LMA under each light condition of one of the cages. Gray asterisks indicate VS collection time window. For DD condition, ambient red light was turned on for 10 minutes. (B) Daily onset (left) averaged (middle) LAM (n cages= 2), presented over days. Each cage had three females and a running wheel for enrichment.

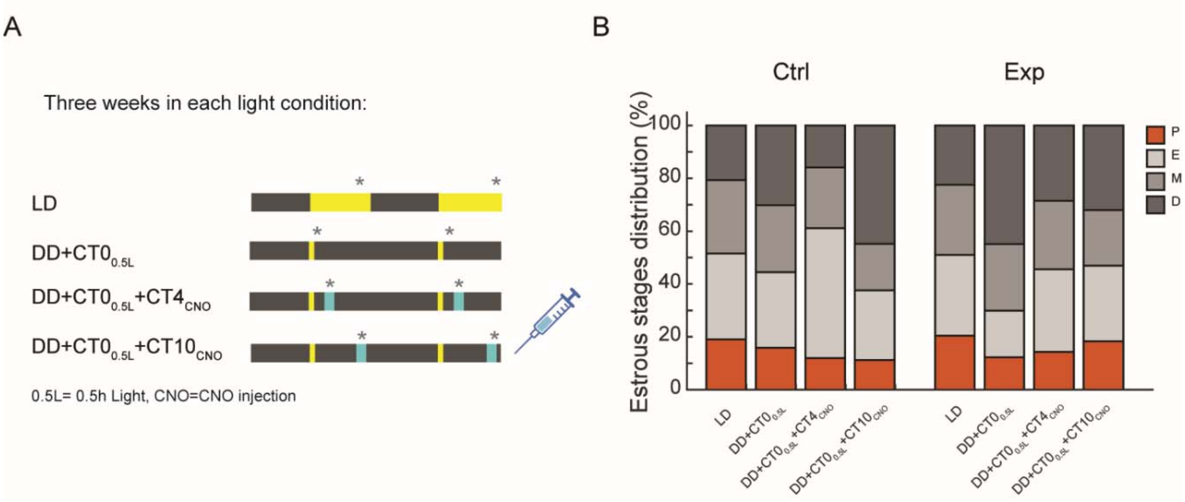


**Figure S8, relate to Figure 5. The estrous-state distributions of $SCN^{VIP}$ chemogenetic activation experiment.** (A) The experimental design (same as Figure 5). (B) Estrous states distributions over three weeks, control (Ctrl, n=6) and experimental (Exp, n=7).

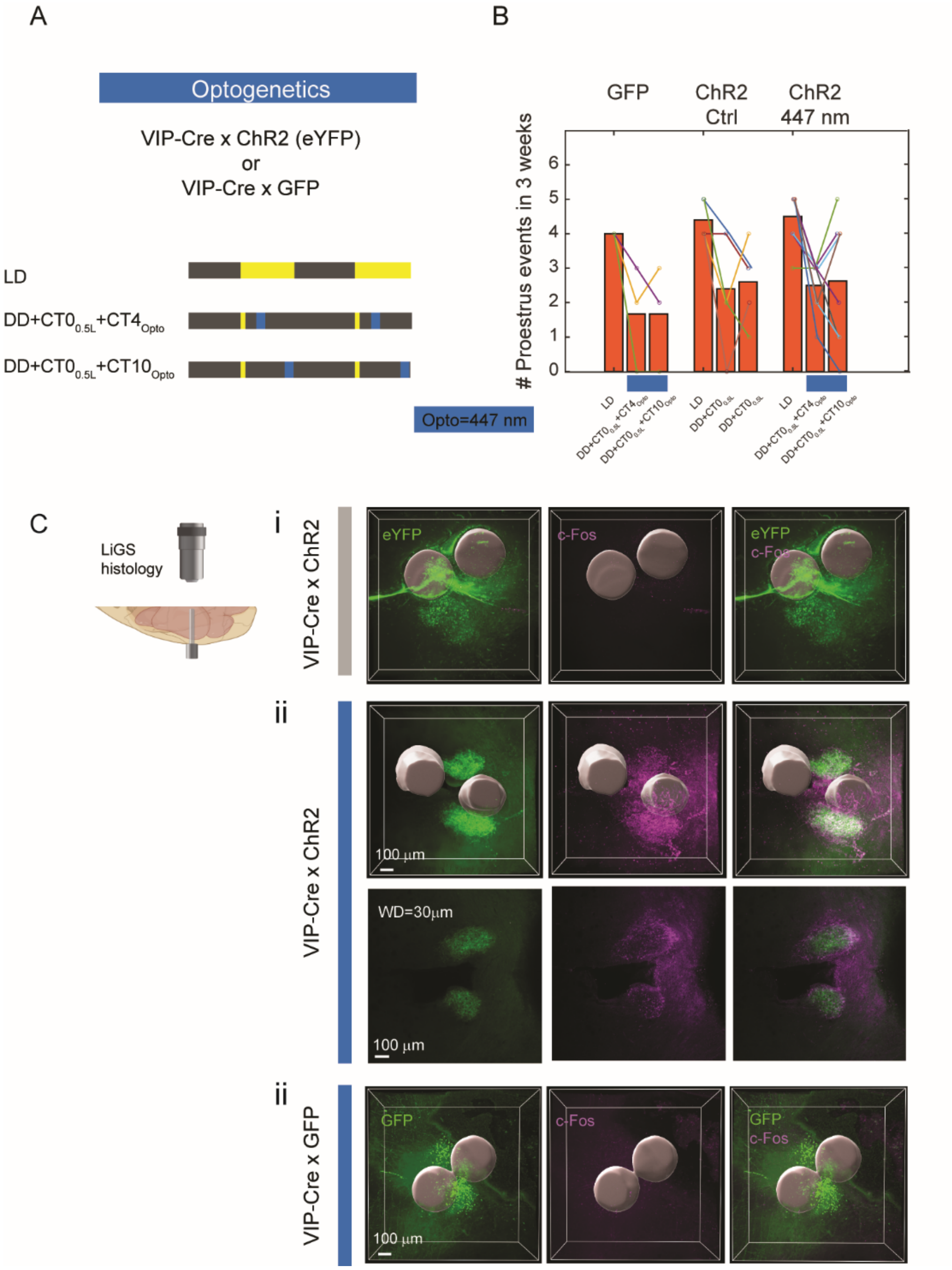


**Figure S9, related to Figure 5. SCN$^{VIP}$ optogenetic activation in the late afternoon is insufficient to rescue estrous cycle regularity.** (A-B) Time-restricted SCN$^{VIP}$ neurons activation with ChR2. (A) The experimental design. VIP-Cre females crossed to ChR2 (ctrl: n=5, 447 nm excitation: n=8) or GFP reporter lines (n=3), put under three conditions, LD, DD+CT0$_L$+CT4$_{opto}$ (447nm excitation at CT4) and DD+CT0$_L$+CT10$_{opto}$ (447 nm excitation at CT10, blue, CT0$_L$ is light for 0.5h at CT0). (B) The number of estrous cycles in three weeks, under the three conditions shown in D. (C) Detailed LiGS histology of SCN$^{VIP}$ neurons below the optical implant. Examples of LiGS histology for *post hoc* fiber location verification and c-Fos expression. (i) VIP-Cre x ChR2 mouse with no neuronal activation. (ii) VIP-Cre x ChR2 mouse with neuronal activation, showing c-Fos expression ~1h after illumination with 447 nm laser, at 3D (top) and 2D (bottom). (iii) VIP-Cre x GFP mouse with c-Fos. Illustration of the fiber in gray, ChR2 or GFP signal, indicative of VIP neurons in green, c-Fos staining ~1h after illumination with 447 nm laser (magenta), showing no expression. 447nm laser illumination was done around ZT14 for 1h.

1. T. Goto, K. Miyamichi, Dynamics of Pulsatile Activities of Arcuate Kisspeptin Neurons in Aging Female Mice. *bioRxiv*, 2022.2008.2008.503241 (2022).
2. A. Cetin, S. Komai, M. Eliava, P. H. Seeburg, P. Osten, Stereotaxic gene delivery in the rodent brain. *Nature Protocols* **1**, 3166-3173 (2006).
3. G. Paxinos, K. B. J. Franklin, *The Mouse Brain in Stereotaxic Coordinates*. (Harvard, 2004).
4. P. Campos, A. E. Herbison, Optogenetic activation of GnRH neurons reveals minimal requirements for pulsatile luteinizing hormone secretion. *Proceedings of the National Academy of Sciences* **111**, 18387-18392 (2014).
5. K. Y. Chan *et al.*, Engineered AAVs for efficient noninvasive gene delivery to the central and peripheral nervous systems. *Nature Neuroscience* **20**, 1172-1179 (2017).
6. D. Gowanlock R. Tervo *et al.*, A Designer AAV Variant Permits Efficient Retrograde Access to Projection Neurons. *Neuron* **92**, 372-382 (2016).
7. R. C. Challis *et al.*, Systemic AAV vectors for widespread and targeted gene delivery in rodents. *Nature Protocols* **14**, 379-414 (2019).
8. J. R. Cho *et al.*, Dorsal Raphe Dopamine Neurons Modulate Arousal and Promote Wakefulness by Salient Stimuli. *Neuron* **94**, 1205-1219.e1208 (2017).
9. Talia N. Lerner *et al.*, Intact-Brain Analyses Reveal Distinct Information Carried by SNc Dopamine Subcircuits. *Cell* **162**, 635-647 (2015).
10. C. K. Kim *et al.*, Simultaneous fast measurement of circuit dynamics at multiple sites across the mammalian brain. *Nature Methods* **13**, 325-328 (2016).
11. A. C. McLean, N. Valenzuela, S. Fai, S. A. L. Bennett, Performing Vaginal Lavage, Crystal Violet Staining, and Vaginal Cytological Evaluation for Mouse Estrous Cycle Staging Identification. e4389 (2012).
12. J. A. McHenry *et al.*, Hormonal gain control of a medial preoptic area social reward circuit. *Nat Neurosci* **20**, 449-458 (2017).
13. L. Brown, S. Hasan, R. Foster, S. Peirson, *COMPASS: Continuous Open Mouse Phenotyping of Activity and Sleep Status [version 1; referees: 3 approved, 1 approved with reservations]*. (2016), vol. 1.
14. K. Gouveia, J. L. Hurst, Improving the practicality of using non-aversive handling methods to reduce background stress and anxiety in laboratory mice. *Scientific Reports* **9**, 20305 (2019).
15. C. Mazuski *et al.*, Entrainment of Circadian Rhythms Depends on Firing Rates and Neuropeptide Release of VIP SCN Neurons. *Neuron* **99**, 555-563.e555 (2018).
16. A. Kahan *et al.*, Light-guided sectioning for precise *in situ* localization and tissue interface analysis for brain-implanted optical fibers and GRIN lenses. *Cell Reports* **36**, (2021).
17. M.-T. Ke, S. Fujimoto, T. Imai, SeeDB: a simple and morphology-preserving optical clearing agent for neuronal circuit reconstruction. **16**, 1154 (2013).
18. P. Berens. (MATLAB Central File Exchange, 2022).
